# Geometric principles underlying the proliferation of a model cell system

**DOI:** 10.1101/843979

**Authors:** Ling Juan Wu, Seoungjun Lee, Sungshic Park, Lucy E. Eland, Anil Wipat, Seamus Holden, Jeff Errington

## Abstract

Wall deficient variants of many bacteria, called L-forms, divide by a simple mechanism that does not depend on the complex FtsZ-based cell division machine. We have used microfluidic systems to probe the growth, chromosome cycle and division mechanism of *Bacillus subtilis* L-forms. The results show that forcing cells into a narrow linear configuration greatly improves the efficiency of cell growth and chromosome segregation. This reinforces the view that L-form division is driven by an excess accumulation of surface area over volume. Cell geometry was also found to play a dominant role in controlling the relative positions and movement of segregating chromosomes. The presence of the nucleoid appears to influence division both via a cell volume effect and by nucleoid occlusion, even in the absence of the FtsZ machine. Overall, our results emphasise the importance of geometric effects for a range of critical cell functions and are of relevance for efforts to develop artificial or minimal cell systems.

## INTRODUCTION

The cell wall is an ancient and highly conserved structure that is almost ubiquitous in the bacterial domain (Errington, 2013). It provides a tough, elastic, protective outer layer around the cell and is largely responsible for the characteristic shapes associated with different forms of bacteria (Egan et al., 2017; Rajagopal and Walker, 2017). The wall is the target for many effective antibiotics, and fragments of the wall are recognised by innate immune receptors (Akira et al., 2006). Its most critical general role lies in osmoregulation, enabling bacterial cells in dilute environments to withstand the turgor pressure generated by the high osmolarity of the cytoplasm (Rojas and Huang, 2017). A large number (~30) of normally essential genes are required for synthesis of the material of the wall, and its spatial regulation during cell growth and division (Errington and Wu, 2017; Zhao et al., 2017).

In the light of the multiplicity of important functions for the wall it is surprising that under certain conditions (isotonic to avoid osmotic lysis) many bacteria, both Gram-positive and Gram-negative, that normally have a cell wall, can thrive in a wall-less state, called the L-form (Allan et al., 2009; Errington et al., 2016). Although L-forms can probably inhabit a range of specialised niches in the environment, they have mainly been studied in the context of their possible role in various chronic diseases and recurrent infections (Domingue and Woody, 1997; Domingue, 2010; Errington et al., 2016).

In previous work with the Gram-positive bacterium *Bacillus subtilis* we have shown that L-form growth requires two types of mutations: one that leads to excess membrane synthesis, and one that counteracts the increased cellular levels of reactive oxygen species (ROS) that occur for reasons that are not fully understood in L-forms (Mercier et al., 2013; Kawai et al., 2015; 2019). Upregulation of membrane synthesis can be achieved directly with mutations affecting the regulation of fatty acid synthesis, or indirectly by inhibiting peptidoglycan precursor synthesis (Mercier et al., 2013). In some bacteria, inhibiting peptidoglycan precursor synthesis alone seems sufficient to enable L-form growth (Mercier et al., 2014). While walled bacteria generally divide by a well-regulated binary fission process, division of L-forms of *B. subtilis* and several other bacteria investigated, occurs through a range of poorly regulated and seemingly haphazard events including membrane blebbing, tubulation, vesiculation and fission. Crucially, these division events occur independent of the normally essential FtsZ-based division machine (Leaver et al., 2009; Mercier et al., 2013; Errington et al., 2016; Studer et al., 2016). Our current model for L-form proliferation assumes that division is driven simply by an imbalance between volume and surface area. Support for this idea comes from the fact that we have been unable to identify mutations in genes required for division, other than those that upregulate membrane synthesis (Mercier et al., 2013). Furthermore, there is a sound mathematical basis for the process (Svetina, 2009) and it has even been replicated in vitro with simple lipid vesicle systems (Peterlin et al., 2009). The simplicity of this division process has led to suggestions that L-form division may be a good model for studying how primordial cells proliferated before the invention of “modern” protein based division machines (Leaver et al., 2009; Chen, 2009; Briers et al., 2012; Errington, 2013). It is also of interest as the basis for proliferation in simplified or artificial cell systems (Blain and Szostak, 2014; Caspi and Dekker, 2014; Hutchison et al., 2016).

Detailed analysis of L-form proliferation has been hampered by the lack of effective systems for following their growth and division by time-lapse imaging. The cells tend not to remain in focus in liquid culture and attempts to tether them to surfaces can cause flattening and lysis. Thus, many questions about their cell cycle remain unresolved, particularly the extent to which chromosome replication and segregation can be controlled and coordinated with growth and division in cells with pleomorphic shape and no cell wall. (Note that in this paper because many of the cells observed are not undergoing division, we use the term segregation for sister chromosomes that have visibly separated, whether or not a division septum separates them.)

Here we report that the use of microfluidic devices that force L-forms into an elongated shape, with cross section similar to that of walled cells, dramatically improves the rate of growth and the efficiency and fidelity of chromosome segregation and other cell cycle processes. The cross section also influences the rate of division in channels. Despite the lack of requirement for FtsZ, division is strongly biased to internucleoid spaces, as in walled cells. Our results also support the notion of a key role for changes in surface area to volume underlying L-form division. Overall, these results show that simple geometric effects can have a profound impact on the efficiency of fundamental cell cycle processes including growth, chromosome replication and cell division. They also lend support to the idea that simple biophysical effects such as phase separation and entropic de-mixing (e.g. Jun and Wright, 2010; Wu et al, 2019b) underlie key steps in the cell cycle of modern bacteria. The results and methods developed here provide important insights into fundamental principles of cell growth, proliferation and chromosome inheritance and have important implications for the development of simplified or artificial cell systems.

## RESULTS

### Irregular division and chromosome segregation in unconstrained L-form cells

Previous work on *B. subtilis* “primary” L-forms (i.e. L-forms derived directly from walled cells, requiring only one or two mutations), as well as many earlier papers with long-propagated “stable” L-forms, have highlighted the inefficient and rather haphazard mode of proliferation in liquid culture (Kandler and Kandler, 1954; Leaver et al., 2009; Studer et al., 2016; Mercier et al., 2013). Fine details of the multiplication process have been difficult to obtain because various methods normally used to fix the position of cells during time-lapse imaging either damage or distort the shape of L-forms, or fail to keep progeny cells in focus. Figure 1A and Movie 1 show typical examples of L-form cells (strain 4740) growing unconstrained in liquid medium in a glass-bottomed microscope dish. Cells clearly underwent growth and division but segments of cell mass frequently moved in and out of focus (top and bottom panels of Figure 1A), making long-term tracking of cells difficult. Note that in this kind of common event, the main cell body appeared to be attached to the glass surface by a fine tube of membrane material which filled up with cytoplasm and DNA as the cell grew. The presence of these fine tubes of membrane has been described in previous L-form publications (e.g. Leaver et al., 2009; Studer et al., 2016), although their nature and biological significance is unclear.

**Figure 1.**
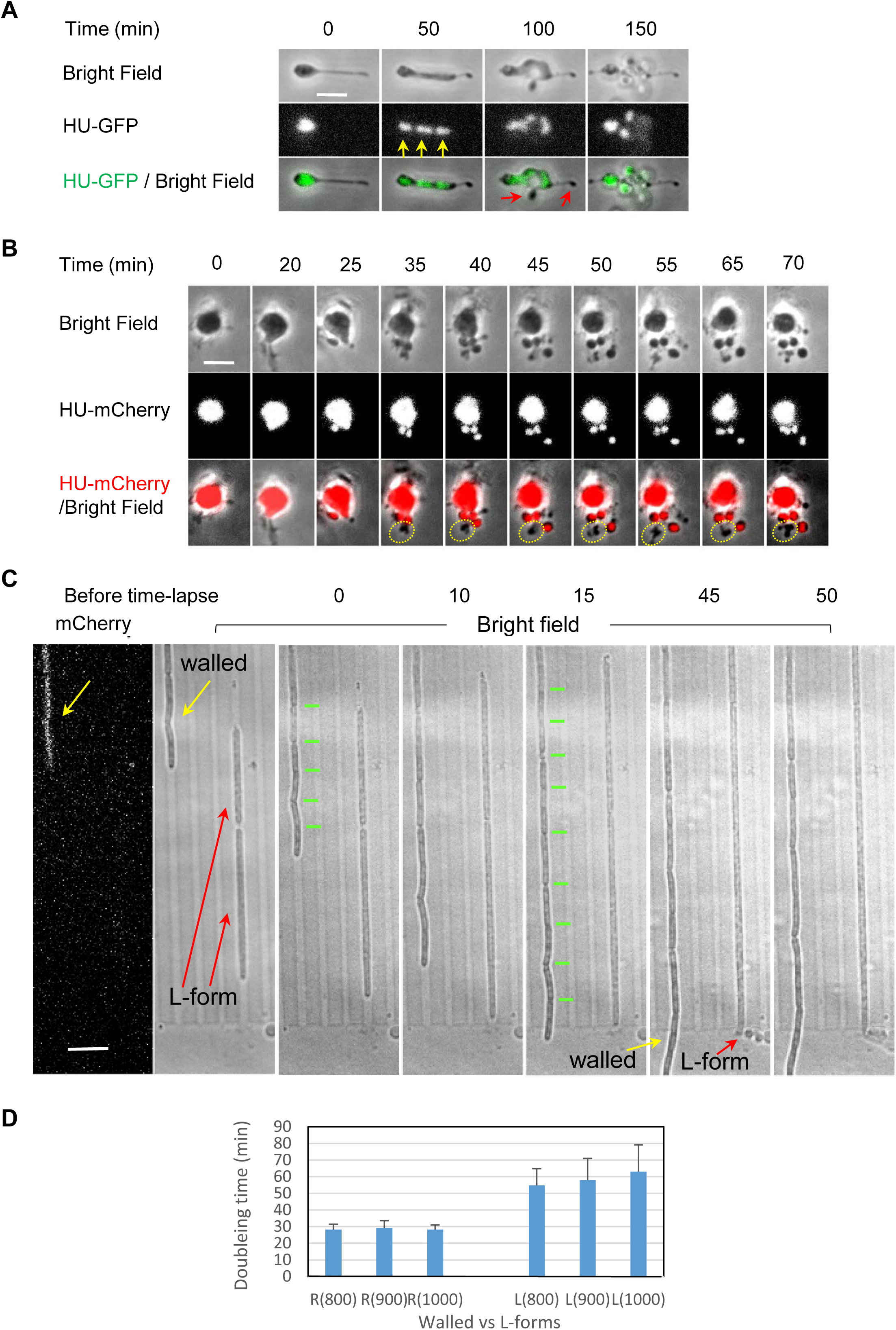
Unconstrained L-form growth in liquid and in microfluidic devices. (A) L-forms growing unrestrained in liquid medium. The figure shows still images from time-lapse microscopy (from Movie 1) of L-form cells of strain 4740 (LR2 *Pspac-dnaA ΩamyE*::*neo hbsU-gfp*) growing in a glass bottomed dish at 30°C. Yellow arrows point to discrete nucleoids and red arrows point to cells devoid of DNA. Scale bar, 5 µm. (B) L-form cells of strain: 4745 (RM121 *ΩamyE*::*hbsU-mCherry rpoC-gfp)* growing in the gutter of a microfluidic system. Example of a series of still images from a time-lapse experiment in which the cells stayed relatively in focus (Movie 2). A daughter bleb with no chromosome is ringed in yellow. For both (A) and (B) each panel shows bright field images on top, chromosomal DNA labelled with HU-mCherry in the middle, and a merge of the two (DNA in green in (A) and red in (B)) at the bottom. The time points (min) of the selected still images from the time-lapse experiment are shown above the images. Scale bar, 5 µm. (C). Growth of walled (strain 4742; LR2 *ΩamyE*::*neo hbsU-gfp aprE::P_rpsD_-mcherry spc ΔxylR::tet*) and L-form (strain 4739; LR2 *ΩamyE*::*neo hbsU-gfp*) *B. subtilis* in microfluidic channels. The pair of images to the left were taken before the time-lapse imaging, showing the mCherry signal in the walled cells on the left (yellow arrows), and two L-form cells that do not express mCherry on the right (red arrows). Short green lines mark division sites. The time points of the selected still images from the time-lapse experiment (5 min intervals) are shown above each image. Scale bar, 5 µm. (D). Growth rates, shown as doubling time (min), of walled and L-form strains in channels of different widths. R: walled cells; L: L-forms. Channel widths (nm) are indicated in brackets.

A key objective of the current work was to characterise the extent to which chromosome replication and segregation remain coordinated with division, so in these and subsequent experiments, chromosomes were labelled with fluorescent fusions to the HU protein, which binds DNA almost non-specifically (Kohler and Marahiel, 1997).

The middle and lower panels of Figure 1A (and Movie 1) showed: (i) that discrete nucleoid-like structures could be discerned within L-forms (yellow arrows) but brighter structures, containing either overlapping multiple nucleoids or un-resolved multiple chromosomes, could be seen frequently (e.g. 0 min); (ii) that these could resolve into multiple discrete structures (e.g. arrows at 50 min); and (iii) that the arrangement in larger L-form clusters was complex and difficult to track because of focus and overlapping problems (e.g. 150 min). It also appeared that some cell lobes might be devoid of DNA (e.g. phase dark objects with no associated fluorescence at 100 min (red arrows). (Note that in this and some subsequent figures the fluorescence image brightness was enhanced to enable visualization of small amounts of DNA. Raw images are available on request.)

### Use of microfluidics to constrain L-form movement during growth

Agarose based microfluidic devices offered a possible way to constrain the cells without damaging them, while maintaining them within a focal plane. We fabricated microfluidic devices based on those described by Moffitt et al. (2012; see Eland et al., 2016; Figure S1). Each device contained sub-micron-scale linear tracks (channels), imprinted into agarose. The channels were restricted in height (~1.6 µm) to impose a strong z-axis control over the cells as they grew. The growth channels were open, at either one or both ends, to gutters through which growth medium flowed, delivering fresh nutrients.

It turned out that the gutters provided an improved way to image L-form growth without physical constraint. Figure 1B & Movie 2 show an example of a common division event within a gutter. Here, a large L-form cell underwent ‘blebbing’ to generate multiple small daughter units, all in fairly good focus. Three of the blebs displayed HU fluorescence in frames from 40 to 70 min, whereas one (circled in yellow in the bottom panels) was non-fluorescent and presumably anucleate. The variation in the sites of division and in the number of nucleoids in daughter cells in these and many other similar experiments showed that chromosome segregation in unconstrained L-forms is poorly regulated and relatively disorganized.

### Imposition of an elongated architecture regularizes L-form growth

Surprisingly, when L-forms were trapped in the channelled area of the microfluidic chamber, so that growth would be forced to occur along a fixed longitudinal axis, a strikingly different pattern of growth was observed. Now, the cells grew rapidly and with uniform appearance along the channel (red arrows in Figure 1C). In the experiment shown, L-forms were mixed with mCherry labelled walled cells (yellow arrows) to enable comparison of their behaviour. The L-forms were almost indistinguishable from the walled cells except that the latter had regular constrictions (due to cell division; indicated by green bars) and a slightly less regular cylindrical shape, perhaps because of frictional drag against the channel walls. However, upon exiting the channels, the difference between walled cells, which continued to grow in straight lines out into the gutter, and the L-forms, which immediately formed chains and clusters of spherical blebs, was striking.

Interestingly, when growing in these channels, the L-forms rarely divided (see the section below). In the typical example shown despite having similar length increase after 15 min of growth in the channels, clear constrictions corresponding to division sites (marked with short green bars) increased from 5 to at least 9 in the chain of walled cells, but none were evident in the L-forms (Figure 1C 15 min).

The microfluidic channel designs in these initial experiments were of two types, featuring repeating patterns of widths approximately 800, 900 and 1000 nm wide, or 600, 700 and 800 nm wide. Walled cells of wild type *B. subtilis* are approximately 850 nm in diameter (Sharpe and Errington, 1998), so the channels roughly mimic walled cell dimensions. Under these conditions the growth rate of the L-forms could be readily estimated from the increase in length over time, assuming that the cross-sectional area of the channel and thus of the cell was constant. As summarised in Figure 1D, in 800 nm channels the average length doubling time of the L-form strain (strain 4739) grown at 32°C was about 2x of that of the isogenic walled cells (strain SL004) (55 min ± 10.1 vs 28 min ± 3.3, respectively).

As expected, growth in the wider channels did not alter the width of walled cells (which normally maintain a constant width irrespective of growth rate; Sharpe et al., 1998), nor did it affect their length doubling time (Figure 1D). The L-forms, however, showed increased length doubling time as the channel width increased (5% for 900 nm and 15% for 1000 nm, respectively).

### Effects of channel width on L-form growth and division

The low frequency of division of L-forms trapped in the narrow channels (Figure 1C, 2A and Figure S2) was unexpected. We previously reported experiments suggesting that L-form division is driven by excess membrane synthesis, creating a high surface area to volume (A/V) ratio that is incompatible with a spherical shape and thus drives shape changes leading to division (Mercier et al., 2013). Cylindrical shapes have a higher A/V ratio than spheres of the same volume. It was therefore possible that the narrow channels imposed a geometry with high enough A/V to eliminate the driving force for division that occurs in unconstrained (roughly spherical) L-forms. If so, increasing the channel width, and therefore reducing the imposed A/V, might re-enable division. To test this we designed two microfluidic chips with wider channels (Chip No. 6 = 1, 1.2 and 1.4 µm; Chip No. 7 = 1.8, 2.0 and 2.2 µm). As predicted, ‘in-channel’ division occurred much more frequently in these wider channels (e.g., 35, 65, 90 and 105 min frames in Figure 2B). It needs to be mentioned that the wide channels were only half the length of those of the narrow channels, and so would effectively give only half the chance of observing ‘in-channel’ division in the same time frame, making direct comparisons difficult. Figure S3A and Movie 3 show a typical example of a long L-form growth sequence in wide channels. Accurate quantitation of division frequency was problematical for several reasons. First, tracking of cells was limited by the channel length, because undivided cells often “bubbled” out of the ends of the channel and this material then disappeared (Figure 1C, 45 & 50 min frames; and 105 min frame onwards in Figure 2A), so measurement of total cell length per division was not possible. Second, after division in the wider channels some progeny cells spontaneously escaped from the channels (e.g. cells labelled with a red star in Figure 2B frames 65 min and 85 min, and the 195 min frame in Figure S3A; Movie 3) so that, again, their subsequent fate could not be recorded. Finally, an element of stochasticity seemed to arise due to small irregularities in the channels, probably either casting irregularities or debris / thin membrane fibres from the growing L-form cells. Nevertheless, we estimated the difference in division frequency by counting ‘in-channel’ division events in continuous cell lineages over 5 hour time courses for channels of different widths. Clustered division events that occurred occasionally in cells with chromosome segregation defects (see below) were excluded from this analysis. The results confirmed that division was rare in narrow (< 1 µm) channels (7 division events in 37 cells during the whole time course) and much more frequent in the wider (1-2.2 µm) channels (72 division events in 38 cell lineages) (Figure 2C). The lower frequency of division in the narrow channels compared with the wide channels is consistent with the A/V model for division in L-forms (see Discussion).

**Figure 2.**
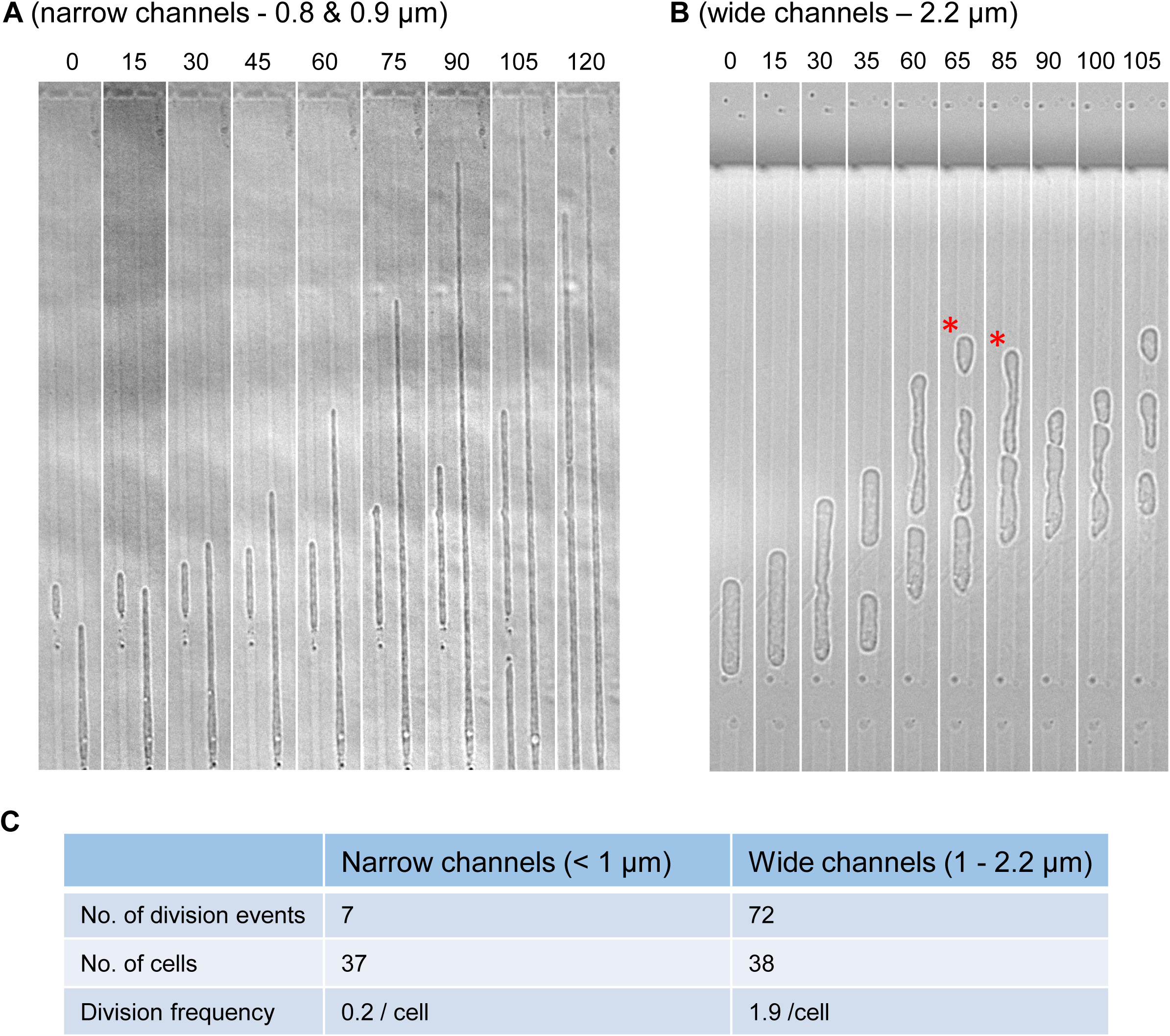
Effect of channel width on L-form division. (A) Lack of division in narrow microfluidic channels. Selected still bright field images of a time-lapse experiment. L-form cells of strain 4739 (LR2 *ΩamyE*::*neo hbsU-gfp*) were loaded into microfluidic chamber (Chip No. 2; channel widths 0.8, 0.9 and 1.0 µm) and grown at 32°C. Images were captured every 5 min. The small cell on the left was in a 0.9 µm channel; the large cell on the right was in a 1.0 µm channel. Full set of still images from this time-lapse series is shown in Figure S2. Scale bar, 5 µm. (B) In wide channels cell division occurred more frequently. Selected still bright field frames from a time-lapse experiment. The cell shown was in the 2.2 µm wide channel (Chip No. 7). Red stars label cells that escaped from the channel. Strain: 4739 (LR2 *ΩamyE*::*neo hbsU-gfp*). Scale bar, 5 µm. (C) Division frequency of L-forms grown in narrow vs wide channels. Only ‘in-channel’ division events were scored. [Tabulated data.]

### Efficient chromosome segregation in channel constrained L-forms

We then examined the effects of channel confinement on nucleoid arrangement and segregation, using HU-GFP fluorescence imaging. When the cells were initially placed in narrow channels, the multiple nucleoids often appeared as large overlapping or un-resolved masses (Figure 3A, red arrows at 0 min; cells in 0.8 and 0.9 µm wide channels) but as the cells increased in length, these masses gradually resolved into smaller, individual nucleoids (e.g. 80 min).

**Figure 3.**
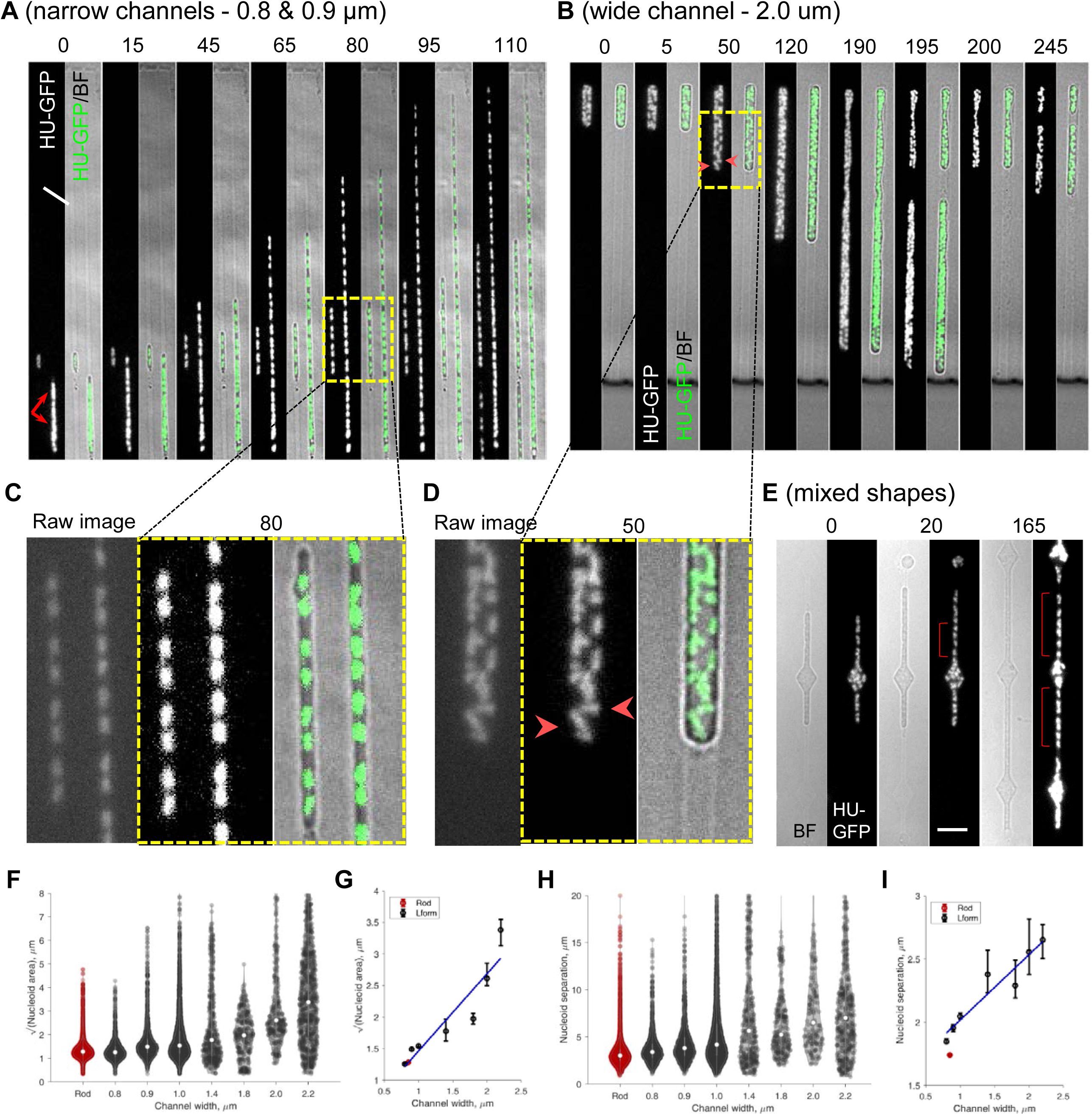
Effect of channel width on nucleoid appearance and arrangement in L-forms. (A, C) Regular chromosome segregation in narrow microfluidic channels. Selected still images of the same time-lapse experiment shown in Figure 2A. For each time frame (time indicated above the images) shown a set of 2 images are presented: a HU-GFP image showing the nucleoids on the left and the merge of the GFP image and the corresponding bright field image on the right. The small cell on the left was in a 0.9 µm channel; the large cell on the right was in a 1.0 µm channel. Full set of still images from this time-lapse series is shown in Figure S2. Yellow boxed region is enlarged and shown in Figure 3C. An un-processed GFP image at t80 min is shown to the left in Figure 3C. Scale bar, 5 µm. (B, D) Irregular chromosome arrangement in wide channels. In wide channels chromosome distribution is irregular. These are selected still frames from Movie 4. Each time frame shows a HU-GFP image showing the nucleoids on the left and the merge of the GFP image and the corresponding bright field image on the right. The cell shown was in a 2.0 µm wide channel (Chip No. 7). Arrowheads: chromosomes lying horizontally or perpendicularly. Strain: 4739 (LR2 *ΩamyE*::*neo hbsU-gfp*). Yellow boxed region is enlarged and shown in Figure 3D. An un-processed GFP image at t50 min is shown to the left in Figure 3D. Scale bar, 5 µm. (E) Chromosome arrangement in Chip No.33 which contained alternating narrow channels and diamond shapes. These are selected still frames from Movie 6. Each time frame shows a bright field image on the left and chromosomal DNA labelled with HU-GFP on the right. Strain: 4741 (LR2 *ΩamyE*::*neo hbsU-gfp aprE::P_rpsD_-mcherry spc)*. Brackets indicate regions where chromosomes appear regularly distributed. Scale bars, 5 µm. (F – I) Quantitative analysis shows that spatial confinement of L-forms can reproduce near-native nucleoid segregation. (F) Nucleoid size, as measured by the square root of nucleoid area, for walled cells (Rod, red) and L-forms in different channel widths (black). (G) Scatter plot of square root of nucleoid area versus channel width for walled cells (red) and L-forms (black). (H) Nucleoid separation for walled cells (Rod, red) and L-forms in different channel widths (black). (I) Scatter plot of nucleoid separation versus channel width. Blue line in (G) & (I): linear fit to the L-form data. For walled cells (red) in (G) & (I), the “channel width” was set to 850 nm to match the known cell width. Violin plots: Circles indicate median, bars indicate upper and lower quartile. Scatter plots: Circles indicate median, error bars indicated 95 % confidence interval from bootstrapping. n=17766 time-lapse observations of nucleoids descended from 45 mother cells in separate agarose channels.

After this initial phase of resolution many cells showed a remarkably regular pattern of chromosome replication and segregation. For example, in Figure 3A (full sequence in Figure S2), over a time frame of 110 min, the short cell on the left, containing one nucleoid at 0 min, undertook three sequential successful duplications (times 15, 65 & 110 min), to give 2, 4 and then 8 segregated nucleoids, while the large cell on the right also showed increasingly regular nucleoid arrangement (times 80 and 95 min) (enlarged section shown in Figure 3C). Movie 4 shows another example of large DNA masses resolving into smaller and often regularly spaced nucleoids. Several L-form strains with different genetic origins were tested (including strains 4739, 4741 and 4744; Table S1) and all were able to resolve large nucleoids and then regularly distribute the chromosomes when grown in narrow channels. Quantitative analysis of various nucleoid parameters (area, width, eccentricity and internucleoid separation; Figure 3F-I & Figure S3D & E), showed that, except for eccentricity (see below), L-form nucleoids in 0.8 or 0.9 µm channels appeared remarkably similar to those of walled cells.

These results demonstrate that cell wall synthesis is not required for regular chromosome segregation, at least not when cells are forced to grow under these geometric constraints. Importantly, these findings also definitively exclude any models for chromosome replication or segregation that require pre-existing markers in the cell wall.

### Effects of cell geometry on chromosome segregation

We then examined the effects of channel width on chromosome replication and segregation. Unlike the narrow channels, chromosome arrangement was increasingly perturbed in the wider L-form cells. Stills of typical frames are shown in Figure 3B, with more examples shown in Figure S3A, B and movies 3 & 5. A close up of the typical nucleoid appearance in a wide channel is shown in Figure 3D. Although nucleoid lobes similar in size and fluorescence intensity to the individual nucleoids of cells in the narrow channels were evident, they tended to form clumps that split up only infrequently. Inspection of the movies revealed highly dynamic patterns of splitting and coalescence that will merit further investigation. Quantitative measurements of various nucleoid parameters relative to channel width are shown in Figure 3 F-I, and Figure S3D & E. Nucleoid area and nucleoid separation (centroid to centroid) both increased in parallel with increasing channel width, due to the failure of nucleoids to separate efficiently in wider channels. The failure of nucleoid lobes to separate was also manifested in a decrease in nucleoid eccentricity (ratio of nucleoid width to length; Figure S3D) and an increase in nucleoid width, which increased proportional to channel width (Figure S3E).

In support of the close connection between cell width and nucleoid configuration, we noticed that when cells grown in wider channels occasionally became slightly constricted, length wise, perhaps because of damage or miscasting of the agarose, nucleoid separation was strikingly improved (red brackets in Figure S3B; Movie 5). To test this further we designed a microfluidic chip with narrow channels interrupted by wider diamond shapes (Figure 3E, S1 & S3C; Movie 6). Nucleoids were well distributed in the narrow (700 nm) part of the channels (red brackets in Figure 3E & S3C; Movie 6). However, on growing into the larger diamond regions, nucleoids lost their regular linear arrangement and spread out in different orientations to fill the space (compare the orientations of the two nucleoids labelled by yellow arrows in Figure S3C, 30 min).

All of these observations and measurements are consistent with the idea that efficient chromosome segregation is dependent on the geometry of the cell and, as is evident from the line plots in Figure 3 G,I, that artificially setting the width of the L-form at about that of walled cells (~850 nm) generates a normal pattern of segregation.

### Division of L-forms mainly occurs between nucleoids

Walled bacterial cells segregate sister chromosomes at cell division with high fidelity. The coordination between segregation and division is thought to rely heavily on an effect called nucleoid occlusion. As first described it was proposed to rely on a phase separation between DNA and cytoplasm, together with a tendency of membrane invagination to be impaired in the nucleoid zone (Valkenburg and Woldringh, 1984; Mulder and Woldringh, 1989; Woldringh et al., 1990). More recently nucleoid occlusion proteins, Noc in *B. subtilis* (Wu and Errington, 2004) and SlmA in *E. coli* (Bernhardt and de Boer, 2005), were identified that are associated with the chromosome and act to inhibit assembly or constriction of the FtsZ machine in its vicinity. Nevertheless, mutants deficient in these proteins still tend to divide between nucleoids under normal conditions (Wu and Errington, 2004; Rodrigues and Harry, 2012). Given that L-form division occurs independently of FtsZ it was interesting to examine whether L-form division is also subject to a nucleoid occlusion effect. Division through the nucleoid is barely detectable in walled cells (Kaimer et al., 2009). Perhaps surprisingly, bisection of nucleoids was also infrequent in L-forms growing in channels. Of 45 division events (excluding the “abnormal” division events that generated anucleate daughter cells – see below) only 4 (9 %) appeared to have occurred through a chromosome (e.g. arrowheads in Figure 4 and Figure S4; Movies 7 & 8). Thus, although the frequency of bisection was much higher than in walled cells using the FtsZ-based division machine, a large majority of division events (91%) still occurred between nucleoids (e.g. arrows in Figure 5A, 105 min).

**Figure 4:**
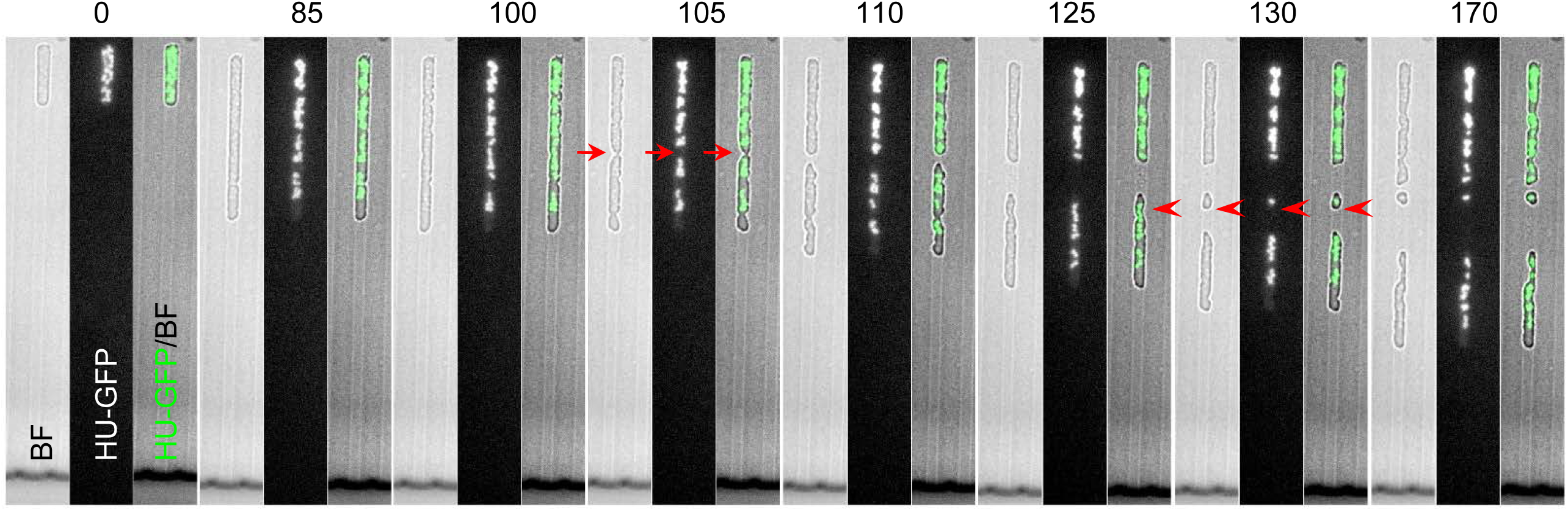
Bisection of chromosomes occurs occasionally in L-forms. A small cell with little DNA (arrowheads in Frame 130 min) appeared to be not growing, possibly because its chromosome is incomplete. These are selected still frames from Movie 7. The cell shown was in a 1.8 µm wide channel (Chip No. 7). Arrows point to division between nucleoids; arrowheads point to possible division through nucleoids.

**Figure 5:**
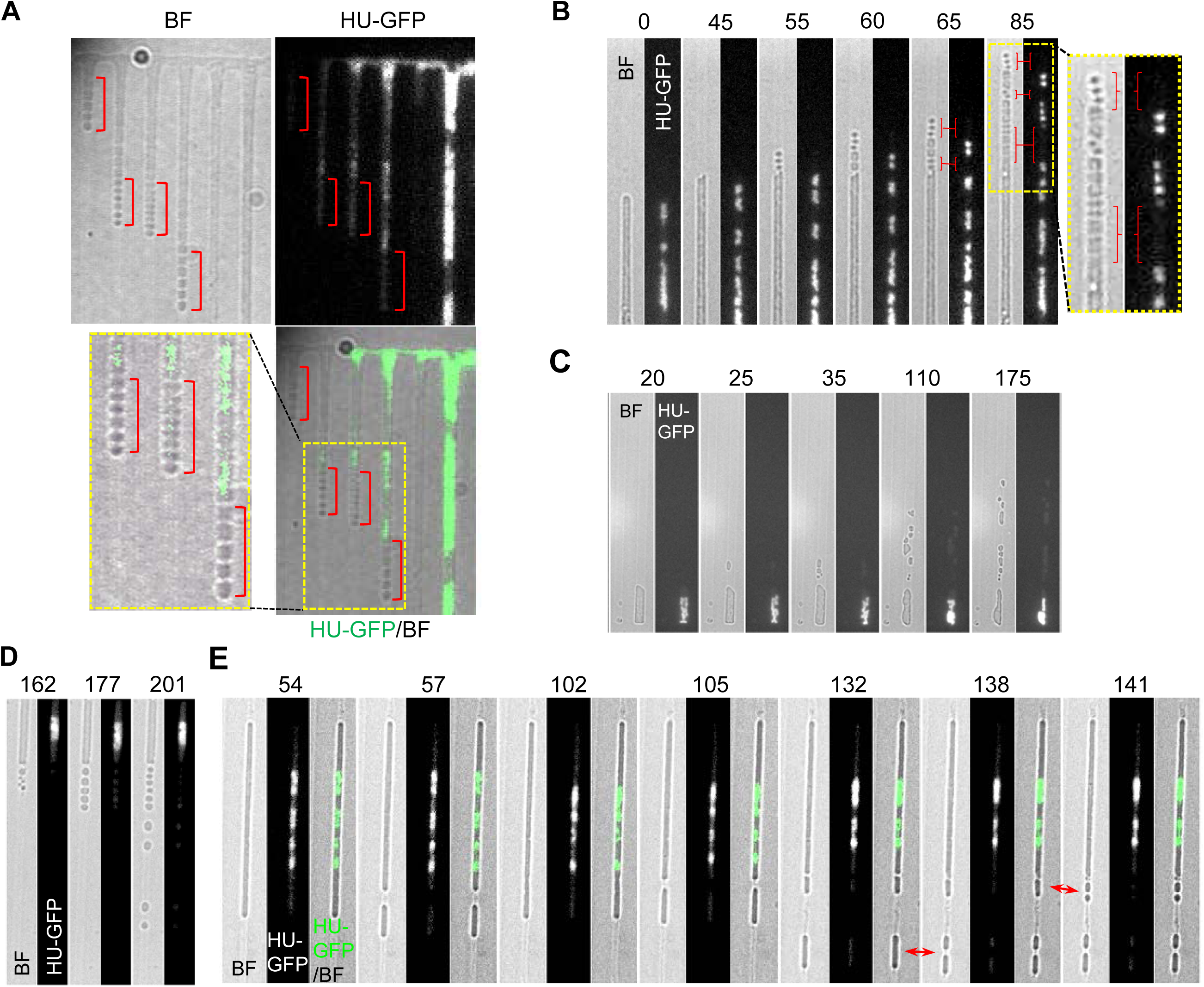
DNA deficiency leads to DNA-free pearling division in channel-confined L-forms. (A and B) DNA-less ‘beads’ produced by L-form in narrow channels under normal growth conditions. Scale bars, 5 µm. Brackets: DNA-free, bead-like cells. (A) Cells in the gutter grew into the narrow channels with the mass of the nucleoid excluded from entry, generating strings of DNA-less beads in narrow channels. A bright field image is shown on the top left, chromosomal DNA labelled with HU-GFP on the top right, and a merge of the two on the bottom right (GFP in green). Yellow boxed region is enlarged and shown at the bottom left. These are selected still frames from Movie 11. Strain 4739 (LR2 *ΩamyE*::*neo hbsU-gfp*). (B) Small DNA-less cells in narrow channels. Each time frame shows bright field images on the left and chromosomal DNA labelled with HU-GFP on the right. Yellow boxed region is enlarged and shown on the left. Strain 4742 (LR2 *ΩamyE*::*neo hbsU-gfp aprE::P_rpsD_-mcherry spc)*. (C) A cell, appeared to be defective in chromosome replication (for unknown reason), produced many small DNA-free daughter cells of various sizes. These are selected still frames from Movie 10. The cell shown was in a 2.2 µm wide channel (Chip No. 7). Each time frame shows bright field images on the left and chromosomal DNA labelled with HU-mCherry on the right. Strain: 4739 (LR2 *ΩamyE*::*neo hbsU-gfp*). Scale bars, 5 µm. (D) and (E) Examples of L-forms inhibited for DNA replication generating regular pearling (D) or large DNA-free cells (E) in the DNA deficient regions in narrow channels. L-forms of strain 4739 (LR2 *ΩamyE*::*neo hbsU-gfp)* were grown in the presence of the DNA replication inhibitor HB-EmAu in liquid culture and after introduction into a microfluidic device. These are selected still frames from Movie 11 (for D) and 12 (for E). Red arrows in (E) shows anucleate cells dividing. Scale bars, 5 µm.

### Division frequency of L-forms is increased by DNA deficiency

We previously postulated that the blebbing or extrusion division events of L-forms could be driven by active nucleoid segregation followed by membrane sealing around the nucleoid (Leaver et al., 2009). This class of model gains support from in vitro experiments showing that encapsulated nanoparticles or macromolecules can drive tubular extrusions or budding transformations in simple lipid vesicles (Yu & Granick, 2009; Terasawa et al., 2012). However, in the channel experiments evidence against this idea arose in rare microfluidic “accidents” of which an example is shown in Figure 5A (full sequence in movie 9). Material spilling over from the filled channel to the right sequentially entered the adjacent channels leftwards. This material appeared to be deficient in DNA presumably because the chromosome entering the channel was incomplete, damaged or delayed. The precise nature of the defect was unclear but it resulted in a striking series of repeated divisions adjacent to the edge of the visible DNA, giving a string of small spherical compartments (red brackets). Figure 5B shows another example in which multiple small anucleate spheres were generated (“pearling”; see Discussion) at the end of a cell and in an unusually large internucleoid region (red brackets in enlarged inset). Similar events occurred in wide channels: the cell in the typical example shown in Figure 5C (and movie 10) appeared to be defective in chromosome replication (no significant change in DNA fluorescence between zero and 175 minutes). Multiple irregular sized anucleate blebs were shed from the cell along the channel.

The above effects appeared to occur generally in cells that were deficient in DNA. To test this idea we set up experiments in which L-forms grown in narrow channels were treated with specific inhibitors of DNA synthesis 6(p-hydroxyphenylazo)-uracil (HPUra) (Brown, 1971) or N3-hydroxybutyl 6-(3-ethyl-4-methylanilino) uracil (HB-EMAU) (Tarantino et al., 1999). The two inhibitors gave similar results, generating elongated cells with few nucleoids, as expected. Importantly, division events were now frequently detected (Figure 5D, E and Movie 11, 12), even in the narrow channels that do not normally support efficient division. Again this always occurred away from regions occupied by a nucleoid. Many cells exhibited pearling (e.g. panel D and Movie 11) but other events were also frequently seen, such as division of the anucleate cells / membrane tubes (Figure 5E, red arrows; Movie 12).

These results appear to exclude the idea that the nucleoid can act positively to promote division and indeed suggest rather that the nucleoid has a negative effect on the division of tubular L-forms.

### Active positioning of nucleoids?

Wu et al. (2019b) showed that single nucleoids in non-dividing *E. coli* cells are robustly positioned at mid-cell, whereas in cells with two nucleoids, they self-organize at 1/4 and 3/4 positions, regardless of the length of the cell. In our experiments with inhibitors of DNA replication, we noticed that some cells with single nucleoids, mainly centrally located (see earlier time frames in Figure 6B and S5C), when divided to generate one DNA-free daughter and one containing the nucleoid, the single nucleoid, which was now asymmetrically located in the cell, moved towards the distal pole to restore its central position (Figure 6 and S5). The movement occurred rapidly, visible within 1 time frame (3 min) after division was observed.

**Figure 6:**
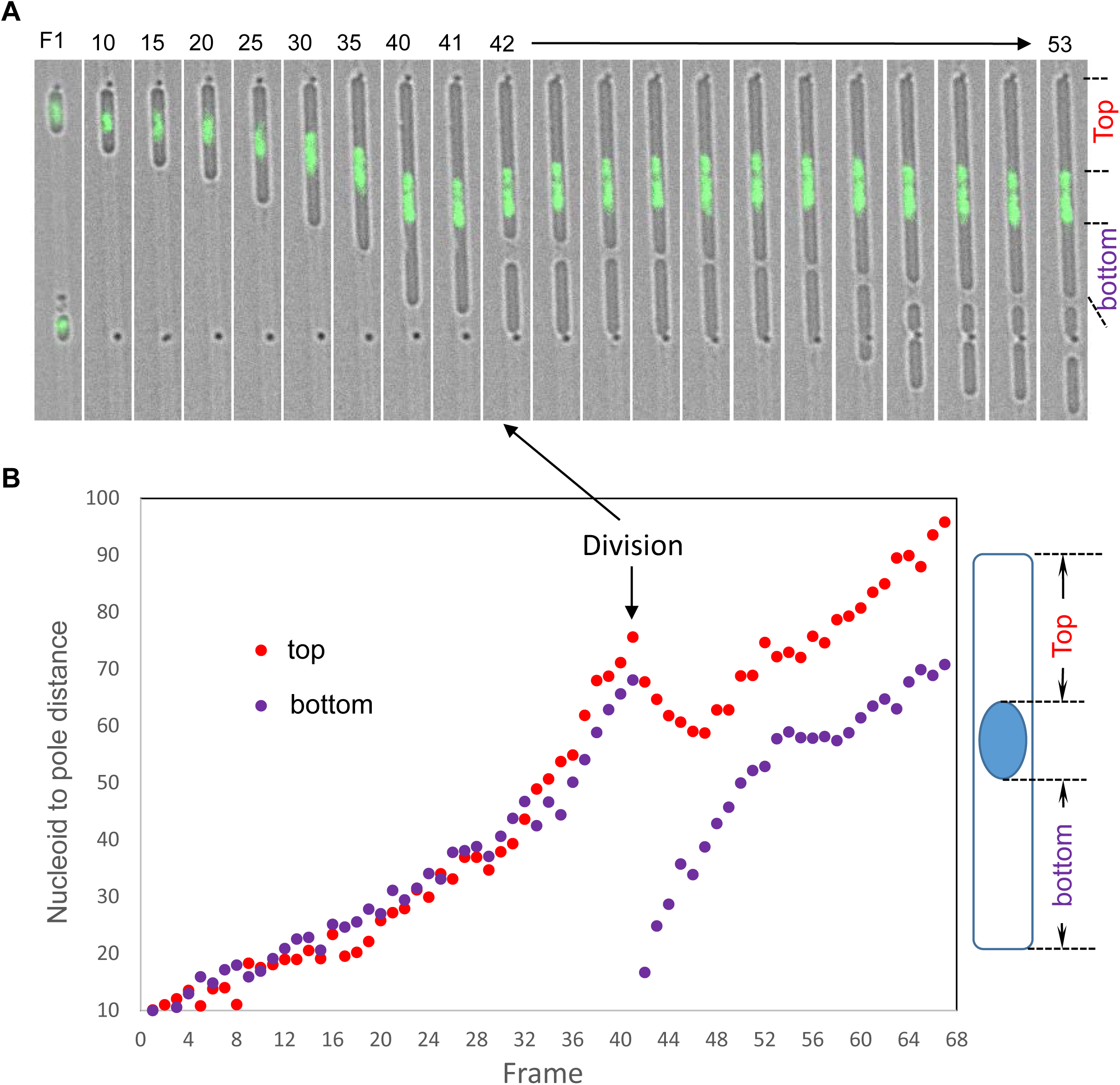
Re-centring of the single nucleoid after asymmetric division in narrow channels. (A) Selected still images showing a single-nucleate cell growing and dividing in the narrow channel. The merge of the bright field image and the green fluorescence image of the chromosomal DNA labelled with HU-GFP (GFP in green) is shown. After division the asymmetrically located nucleoid moved towards the distal pole to re-centre itself. L-forms of strain 4739 (LR2 *ΩamyE*::*neo hbsU-gfp)* were grown in the presence of the DNA replication inhibitor HB-EmAu in liquid culture and after introduction into a microfluidic device. 3 min per frame. Scale bars, 5 µm. (B) Distances of the nucleoid to the two cell poles over a 2.5 h time course. The distance (arbitrary unit) between the polar edge of the nucleoid to the nearest pole was measured. Division that occurred at Frame 42 is marked by arrows.

## DISCUSSION

### Importance of being rod shaped

There is a long literature on L-forms (reviewed by Allan et al., 2009; Errington et al., 2016), including a range of papers describing their irregular almost haphazard mode of proliferation (e.g. Kandler and Kandler, 1954; Gumpert and Taubeneck, 1983; Leaver et al., 2009; Studer et al., 2016). Parents and daughter L-form cells vary greatly in terms of their size, in contrast to the relatively tight ~2-fold variation in the size of most walled bacteria. Division is also quite difficult to define because cells can form blebs or tubes that can retract and re-fuse with the parent cell (Leaver et al., 2009; Studer et al., 2016). However, by forcing L-forms into an elongated configuration similar to that of walled cells, growth and chromosome segregation were all greatly improved. This suggests that geometry plays an important role in cell fitness. Interestingly, Hussain et al (2018) has also observed improved growth for mutant walled *B. subtilis* cells when transitioned from spherical to rod shape, achieved by adjusting the expression level of *tagO* (involved in wall teichoic acid synthesis) and My^2+^ concentration, and proposed that lower doubling time of rods is likely due to cell shape and not another effect. Our result with L-forms clearly excluded the involvement of the cell wall. It thus seems that cell function is tightly connected to geometry because parameters such as surface area to volume ratio, cytoplasm to nucleoid, DNA to protein, membrane to cytoplasm, etc, are all directly affected by cell geometry.

Why should an axial organization, as imposed by the channel or a cylindrical wall be a preferred state? The subject has been reviewed in detail by Young (2006). Obvious possibilities include the following. First, a higher A/V ratio (i.e., a rod shape) can improve nutrient uptake by providing more surface area through which diffusion can occur, and / or more receptors for uptake of specific nutrients. Second, elongation with constant perimeter (i.e. in a cylinder) provides a way to balance the rates of synthesis of cytoplasm and surface: each increment in length (*l*) results in a requirement for 2*π.r.l* in surface area and *π.r^2^.l* in volume. If the radius (*r*) is constant, as would normally be the case, cell surface area is directly proportional to volume, irrespective of length. Third, the geometry provides cells with an axis of polarity along which the segregation of chromosomes can occur.

### Mechanism of L-form proliferation

L-form proliferation has recently been identified as an interesting paradigm for how primitive cells could have proliferated before the invention of the cell wall, and as an interesting starting point for the development of artificial or minimal cell systems (Leaver et al., 2009; Briers et al., 2012; Errington, 2013; Blain and Szostak, 2014; Caspi and Dekker, 2014; Hutchison et al., 2016). Genetic experiments showed that various mutations enabling the proliferation of *B. subtilis* L-forms all had in common an upregulation of cell membrane synthesis (Mercier et al., 2013), leading to the suggestion that excess surface area synthesis could drive proliferation. Theoretical considerations backed up by simple in vitro systems have demonstrated how an increase in surface area at constant volume can drive simple membrane vesicles to divide (e.g. Kas & Sackman, 1991; Peterlin et al., 2009). In rod shaped bacteria, cell surface area (A) increases almost proportionately to cell volume (V). However, in spherical cells the A/V ratio decreases during growth. It is not clear what happens to surface area regulation when normally rod-shaped bacteria transition to a non-rod shape, e.g. when the *mreB* system is impaired or the cell wall is lost. Bendezu & de Boer (2008) showed that *E. coli* cells transiting to a spherical (but walled) mutant state accumulate intracellular vesicles that presumably accommodate excess surface material, indicating that *E. coli* tends not to downregulate surface synthesis under these circumstances. However, *B. subtilis* seems to be able to regulate membrane synthesis, as spherical (*rod*) mutants do not seem to generate intracellular vesicles and, as mentioned above, *B. subtilis* L-forms require upregulation of membrane synthesis to proliferate (Mercier et al., 2013).

Although it will clearly be interesting to follow up with a more detailed quantitative analysis it is apparent that the narrow channels, which impose a higher A/V ratio on the L-forms, almost eliminate division, whereas the wider channels allow division to occur frequently. These observations are consistent with our previous model for L-form proliferation (Mercier et al., 2013) in which division is driven by the rate of surface growth exceeding that of volume increase. Shape changes and ultimately the formation of several smaller progeny from one large L-form dissipate the excess surface area generated during growth.

### Effects of cell geometry on chromosome positioning and segregation

Chromosome spacing and orientation were strikingly improved by the confinement of L-forms in narrow channels. An important conclusion from these experiments is that they exclude any models for chromosome segregation that invoke an essential requirement for specific interactions with the cell wall or cell poles.

Several well conserved proteins have been implicated in chromosome organization and or movement. The ParAB proteins play a reasonably well defined role in segregation of low copy number plasmids, and homologues are found in the chromosomes of most bacteria. Work on *Caulobacter* and sporulating cells of *B. subtilis* (Wu and Errington, 2003; Toro et al, 2008; Shebelut et al 2010; Ptacin et al, 2010; Lim et al, 2014; Wang, 2014; Kloosterman) has revealed that these proteins have an active role in movement of origin regions towards cell poles. However, *Caulobacter* is unusual in having highly specialized cell poles and work here and in filamentous *B. subtilis* suggests that poles are not important outside of sporulation. The SMC or MukBEF protein complexes are also found in virtually all bacteria and appear to work by helping to self-condense chromosomes, inhibiting the formation of tangles.

However, how sister chromosomes come to occupy different spaces and ultimately move away from each other remains poorly understood in bacteria. Much recent work has focused on the role of entropic forces to drive segregation (Jun and Mulder, 2006; Jun and Wright, 2006; Mondal et al., 2011; Minina and Arnold, 2014; Shi and Huang, 2019). Dekker and colleagues have shown that chromosome size, configuration and positions are markedly influenced by the geometry of confinement in non-dividing walled cells perturbed in various ways (Wu et al., 2019a; 2019b). The behaviour of L-form nucleoids in our channel experiments generally support their findings. Importantly, our surprising observation of active and rapid re-centring of nucleoids after division further highlights the importance of biophysical effects in bacterial chromosome positioning and segregation.

### Possible influence of the nucleoid in L-form division

Several theoretical and practical papers have highlighted the possible role of macromolecules or nanoparticles in promoting the division of cells or vesicles (e.g. Yu and Granick, 2009; Terasawa et al., 2012). Based on these ideas it seemed possible that segregating nucleoids could drive the proliferation of L-forms by acting as nanoparticles. This would have an important knock-on effect in that it would ensure that progeny L-form cells contain at least one chromosome. However, our channel experiments showed that involvement of the nucleoid in L-form division is complex. First, our results provided further support for the old idea of “nucleoid occlusion”, in which the nucleoid has a localized negative effect on cell division, thereby helping to ensure that progeny cells have intact chromosomes (Mulder and Woldringh, 1989; Woldringh et al., 1991). While the recent identification of protein factors, such as Noc in *B. subtilis* and SlmA in *Escherichia coli*, provided the first molecular mechanisms for the nucleoid occlusion effect, it is also clear that in both organisms, the cell division machine is still biased away from the nucleoid in the absence of Noc or SlmA (Wu and Errington, 2004; Bernhardt and de Boer, 2005; Wu and Errington, 2012; Moriya et al., 2010). Importantly, our results clearly show that this bias also occurs for the FtsZ-independent division of L-forms, perhaps pointing to a more fundamental, biophysical basis for the effect, perhaps based on separation of nucleoid and cytoplasmic phases, or a change in the property of the membrane proximal to the nucleoid due to co-transcriptional translation and translocation (transertion) of membrane and secreted proteins.

The second apparent effect of the nucleoid was more surprising. In cell tubes in which replication or segregation was blocked, either spontaneously or by an inhibitor, chains of small anucleate cells were frequently generated by repeated sequential divisions adjacent to a nucleoid area, even in the narrow channels that do not normally support L-form division. This dramatic effect is reminiscent of a biophysical effect called pearling instability, which occurs when the membrane material in cylindrical lipid tubes is subject to tension, e.g. by the action of laser tweezers (Bar-Ziv and Moses, 1994; Nelson et al., 1995; Chen, 2009), or the budding transition of phospholipid vesicles that leads to the formation of a chain of vesicles at the increase of the area-to-volume ratio (<u>Käs</u> and <u>Sackmann</u>, 1991). It seems possible that in L-forms these events occur because DNA normally contributes a large proportion of the total mass or effective volume of the cytoplasm. Assuming that synthesis of membrane and cytoplasm (all cell contents other than the nucleoid mass) continue at their normal rates the loss of nucleoid expansion would give an overall increase in A/V, which we have shown previously to drive L-form proliferation (Mercier et al., 2013). That the pearling events always occurred in regions deficient in or devoid of DNA provides further strong evidence that nucleoids inhibit division. It is interesting to note that the possible role of DNA in cell volume regulation has been highlighted recently by experiments analysing the inflation of the *B. subtilis* forespore compartment by DNA import (Javier-Lopez et al., 2018).

Finally, our findings have important implications for attempts to use L-form like division in the development of artificial cells. As well as providing further insights into the key parameters that need to be controlled to drive division, they also suggest that rates of DNA, cytoplasm and membrane synthesis all need to be properly balanced to efficiently coordinate division and chromosome segregation.

## METHODS

### Bacterial Strains, Plasmids, and Growth Conditions

The bacterial strains and plasmid constructs used in this study are shown in Table S1. *B. subtilis* transformation was performed by the two-step starvation procedure as previously described (Anagnostopoulos and Spizizen, 1961; Hamoen et al., 2002).

Walled *B. subtilis* cells were grown on nutrient agar (NA, Oxoid) or in Luria-Bertani broth (LB). *B. subtilis* L-forms were grown in osmoprotective liquid medium NB/MSM at 30°C without shaking. The NB/MSM medium is composed of 2x magnesium-sucrose-maleic acid (MSM) pH7 (40 mM MgCl2, 1 M sucrose, and 40 mM maleic acid) mixed 1:1 with 2 x NB (nutrient broth). 0.8% xylose and 0.8 mM IPTG (Isopropyl β-D-1-thiogalactopyranoside) were added as needed. When necessary, antibiotics were added to media at the following concentrations: 5 µg/ml chloramphenicol, 5 µg/ml kanamycin; 55 µg/ml spectinomycin; 10 µg/ml tetracycline; 1 µg/ml erythromycin; 25 µg/ml lincomycin and 200 μg/ml Penicillin G. L-form strains derived from LR2 were maintained in the L-form state by the addition of Penicillin G and omission of xylose in the growth medium, while those derived from RM121 were stable L-forms and did not require the addition of Penicillin G.

### Protoplast and L-form preparation in Liquid medium

Exponentially growing *B*. *subtilis* walled cells (OD600nm of 0.2~0.3) in LB medium with appropriate supplements were harvested and washed once in LB, then resuspended in NB/MSM containing lysozyme (2 mg/ml). The cells were incubated at 37°C with gentle shaking for 1 hr, or until all the rod-shaped cells have been protoplasted. The protoplasts were then diluted (1 in 5000) in NB/MSM containing 200 μg/ml PenG, and grown at 30°C without shaking for 2 to 4 days, during which time the protoplasts would transit into L-forms. The freshly generated L-forms were diluted at least twice in the same medium and cultured at 30°C, before being used for further experiments.

### Microfluidic system and microscopy

Microfluidic experiments were carried out using a device produced in-house based on that described by Moffitt et al (Moffitt et al., 2012; Eland et al., 2016; Figure S1). Each microfluidic design (chip) contains a set of 3 tracks of different widths, repeated and grouped into 15 μm x 20 μm blocks divided by gutters. The channel widths for the agarose microfluidic chips used are: Chip No.2 (previously: 0.8, 0.9 and 1.0 µm; new: 0.6, 0.7 and 0.8 µm); Chip No.6 (1, 1.2 and 1.4 µm); Chip No. 7 (1.8, 2.0 and 2.2 µm) and Chip No.33 (0.6, 0.7 and 0.8 µm interrupted by diamond shapes). The diamond shapes in Chip No.33 were spaced 20 μm apart (measured from centre to centre) and measure 3.4 μm at the widest point. The designs were initially created on L-edit software and transferred onto a silicon wafer using lithography and deep reactive-ion etching (DRIE), followed by backfilling with a coating of Tetraethyl orthosilicate oxide (TEOS oxide) to produce channels of the desired dimensions (Lionex Ltd). Using TEOS oxide backfill negates the need to use expensive e-beam lithography. Replica moulding of the silicon wafer with hard PDMS was used to create the intermediate mould used to transfer the pattern onto the agarose. Patterned agarose pads, with channels ~1.6 μm high, were cast using an intermediate PDMS mould and an ‘agarose casting mould’ made of PDMS, with 4% low melting point agarose (SeaPlague GTG Agarose from Lonsza, gelling temp 26–30°C) in growth medium containing 1x MSM and 1/5x NB, which then set slowly at 30°C for 1 – 2h.

The structural part of the device, the PDMS chamber block, was cast using a custom designed and milled aluminium mould that matches the size of the mould for casting the agarose pad. Plasma-bonding of the PDMS chamber block to a long cover glass (Agar Scientific Ltd, L4239-2, Coverglass 35×64 mm No.1.5) created a sample chamber. The cover glass which formed the bottom of the sample chamber was coated with BSA (0.5 mg/ml) and allowed to dry. 5 μl of concentrated L-form culture was added onto the cover glass in the sample chamber, then the patterned agarose pad placed (patterned side down) onto the cells in the sample chamber, trapping bacterial cells in the channels of the agarose pad. The chamber was then sealed with a plasma-treated cover glass (Agar Acientific Ltd, L46s20-5, coverglass 20×20 mm No.5) to the top of the agarose pad. The assembly was left at 30°C for 20 min to allow plasma bond to set. The PDMS chamber block also contained two buffer reservoirs on either side of, and connected to, the sample chamber, one for imputing fresh medium and the other as the outlet of the spent medium and bacterial cells that were not confined in the tracks. Modified growth medium with 1/5x strength of NB was supplied continuously through the inlet reservoir from a 50 ml syringe, controlled by a syringe pump at a speed of 1 ml/h using the WinPump Term software (New Era).

Microscopy was performed on a Nikon Eclipse Ti inverted fluorescence microscope System, fitted with an Apo TIRF objective (Nikon 60×/1.49 Oil), as described previously ((Kawai et al., 2015)). All time-lapse experiments were carried out at 32°C unless otherwise indicated. For experiments with the DNA replication inhibitor HB-EMAu (N3-hydroxybutyl 6-(3′-ethyl-4′-methylanilino) uracil; Tarantino et al., 1999), freshly growing L-form culture was mixed with the inhibitor at 3 µg/ml then loaded into the microfluidic devices. Sometimes the mixture was incubated at 30°C from 30 to 40 min prior to loading. The inhibitor was also added to the flow medium at the same concentration to maintain the inhibition.

Time-lapse microscopy of L-forms growing in liquid medium was performed using ibiTreat, 35 mm sterile glass bottom microwell, on a DeltaVision^®^RT microscope (Applied Precision, Washington, USA) as described by Domínguez-Cuevas et al. (Domínguez-Cuevas et al., 2012). Briefly, 200 µl of L-form cells were placed in the dish and leave to stand for 10 min. To adhere the cells to the surface of the glass dish, the dish was centrifuged at 100 g for 5 min using a Beckman Allegra X-12R centrifuge.

### Quantitative image analysis

Movies were prepared for quantitative analysis in the following steps. Images were registered to correct for drift using FIJI/ ImageJ StackReg (Schindelin et al, 2012). Images were manually rotated such that the microchannels were precisely vertical (FIJI, bicubic interpolation). Images were background subtracted using FIJI paraboloid rolling ball, radius 50. Cell in each agarose channel were then manually quality controlled to exclude channels initially loaded with more than one cell, or cells where overgrowth from adjacent channels obscured the cell growth in that channel. Cropped data from individual quality controlled channels were then exported for quantitative analysis in MATLAB.

Nucleoids from cells in each channel were segmented using Otsu’s method. Spurs/ connecting noise pixels were removed using image opening with a disk radius 1. Shape and size parameters for segmented nucleoids were then calculated. Cell size was measured by square root of area rather than area because it is a linear quantity. Nucleoid separation was measured as the distance between the centroids of vertically adjacent nucleoids. Nucleoid angle was measured as the angle between vertically adjacent nucleoids, defined such that the angle between two vertically aligned cells was zero degrees. Nucleoid eccentricity was estimated as e=(minor axis length)/(major axis length) of the binary object.

Confidence intervals in Figure 3G, I were estimated by bootstrapping.

Due to the large size of the dataset (*n=17766 nucleoids)* outliers or low frequency extrema were observed in the data. In order to visualise average trends in the data, in Figures 3F, H and S3D, E, it was necessary to zoom in to less than the full data range. The full extent of the data are shown in Figure S6B-E.

## ACKNOWLEDGEMENTS

We would like to thank Drs Neal Brown and George Wright for the gift of HB-EMAu, and Dr J-W Veening for the HU-mCherry and HU-GFP strains. We also thank the Moffitt lab for help and advice on the agarose-based microfluidic system, and John Hedley and Neil Keegan for their advice on the manufacturing of microfluidic moulds. Work in the Errington lab was funded by a Wellcome Investigator Award (209500) and a European Research Council Award (670980). Work in the Wipat lab was funded by grants from the UK EPSRC (grants EP/N031962/1, EP/K039083/1 and EP/J02175X/1).

## AUTHOR CONTRIBUTIONS

L.J.W. and J.E. designed the study. L.J.W. did most of the experiments. S.L. carried out the cell growth rate measurements. S.P., L.E. and A.W. contributed to establishment and development of the microfluidic systems. SH and L.J.W. carried out quantitative image analysis. L.J.W, S.H. and J.E. analysed data and drafted the manuscript.

## SUPPLEMENTAL FIGURE AND VIDEO LEGENDS

**Figure S1.**
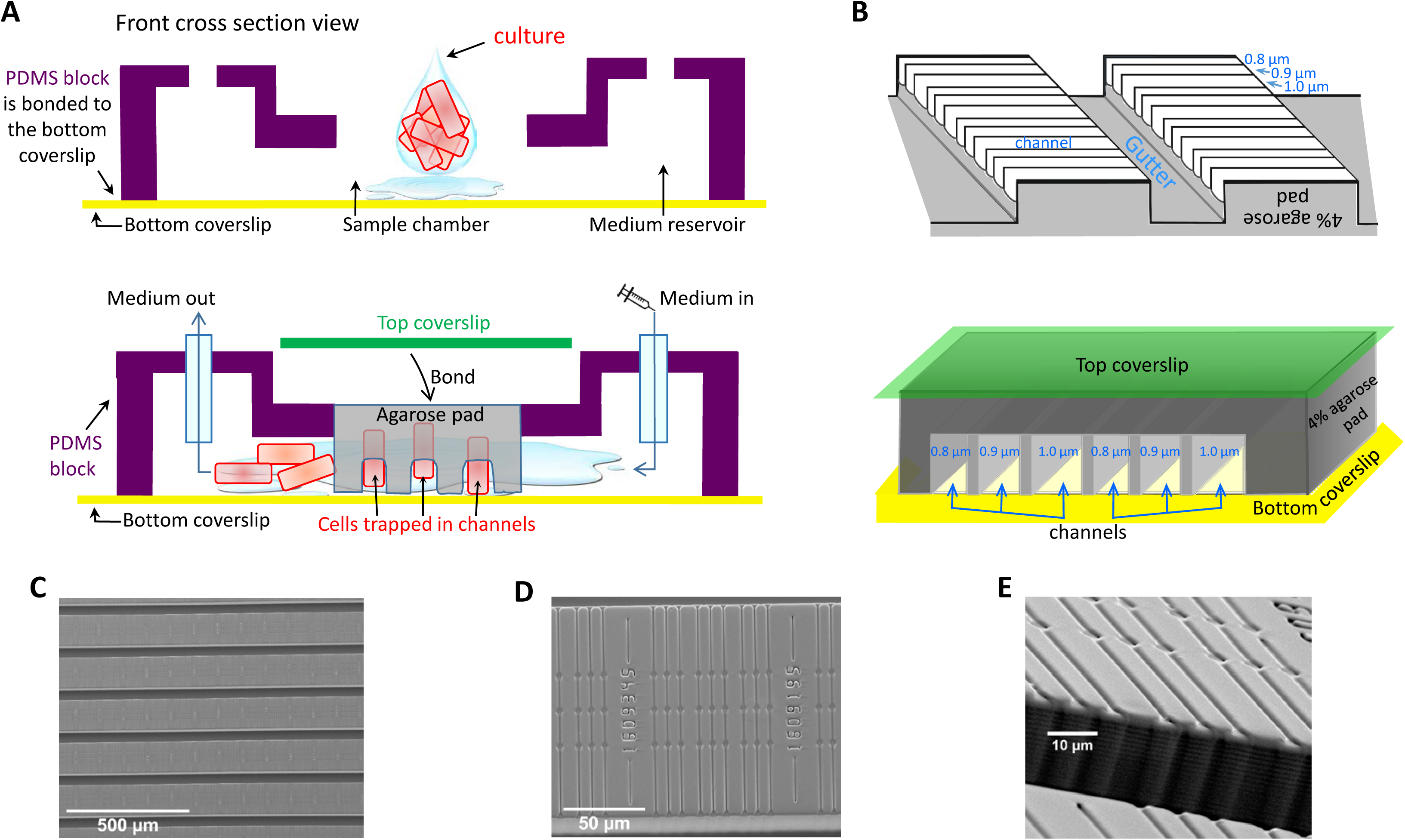
The agarose-based microfluidic device adapted from that of Moffitt et al (2012). Related to Figures 2 to 6. (A, B): Schematic diagrams of the microfluidic system and agarose chips (not to scale). (A) Top: Front cross section view of a partially assembled device. Bonding of the PDMS block to the bottom cover glass creates a chamber ready for cells and the printed agarose pad. The chamber is connected to the reservoirs on the two sides. Bottom: An additional cover glass is laid over the agarose pad, and compresses and seals the device. Cells are confined between the patterned agarose pad and the glass bottom coverslip. Medium flows pass both ends of the channels and removes cells as they emerge from the channels. (B) Top panel: a section of an agarose chip shown here upside-down before being bonded to a cover glass. Each chip consists of sections of tracks. Each section of tracks is ~100×100 µm, containing repeats of a set of three tracks of slightly different widths (0.8, 0.9 and 1.0 µm, for example) and are grouped into 15μm x 20μm blocks divided by gutters. After being mounted onto a cover glass (bottom panel), the glass forms the bottom of the channel. The channel has agarose as the sides and the top, and is open on one end or both ends to the gutter. (C, D, E) SEM images of the surface pattern that is replica moulded onto the agarose pad using the intermediate mould. As agarose cannot be easily imaged with SEM, PDMS was moulded against the intermediate and cured to check the dimensions and structure of the surface pattern. (C) A zoomed out view to show the repeating pattern of the modules, each individually numbered so locations can be monitored. The darker stripes running left to right are the gutters, through which medium flows. The gutters are 40 µm wide. (D) The repeating unit, this image shows Chip No.33. (E) A tilted view of Chip No.33 to highlight the depth of the channel features (1.6 µm) compared to that of the gutters (40 µm).

**Figure S2:**
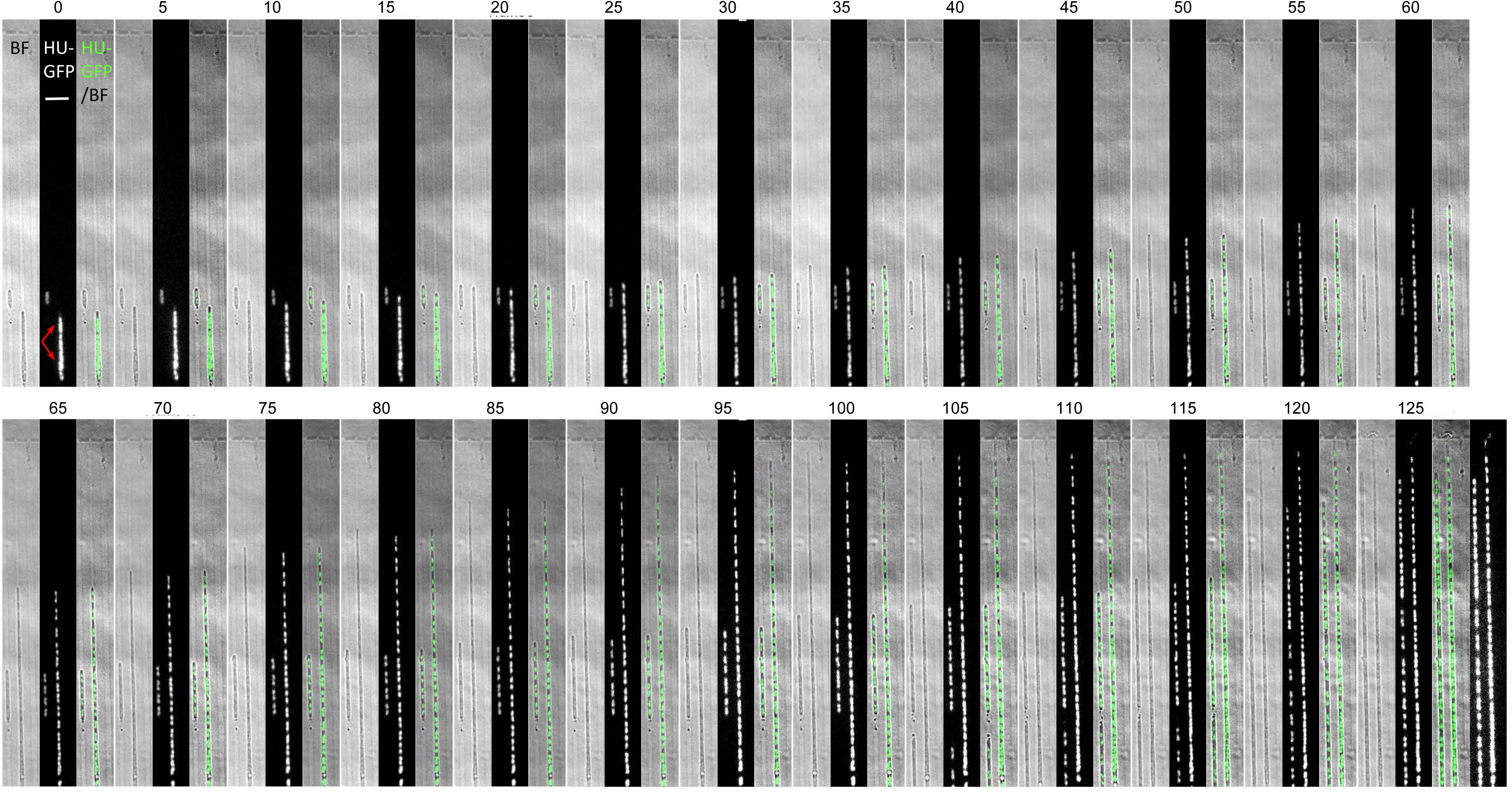
Regular chromosome segregation in narrow microfluidic channels. Related to Figure 2A, 3A and 3C. Full set of still images of a time-lapse experiment presented in Figure 2A and 3A. L-form cells of strain 4739 (LR2 *ΩamyE*::*neo hbsU-gfp*) were loaded into microfluidic chamber (Chip No. 2; channel widths 0.8, 0.9 and 1.0 µm) and grown at 32°C. Images were captured every 5 min. For each frame shown a set of 3 images are presented: a bright field images on the left, a HU-GFP image showing the nucleoids in the middle and the merge on the left. For the last time frame (Frame 26) an extra image of the nucleoids, with increased brightness to show the DNA in cells exiting the channel, is shown to the right. The small cell on the left was in a 0.9 µm channel; the large cell on the right was in a 1.0 µm channel Arrows: un-resolved chromosomal mass.

**Figure S3:**
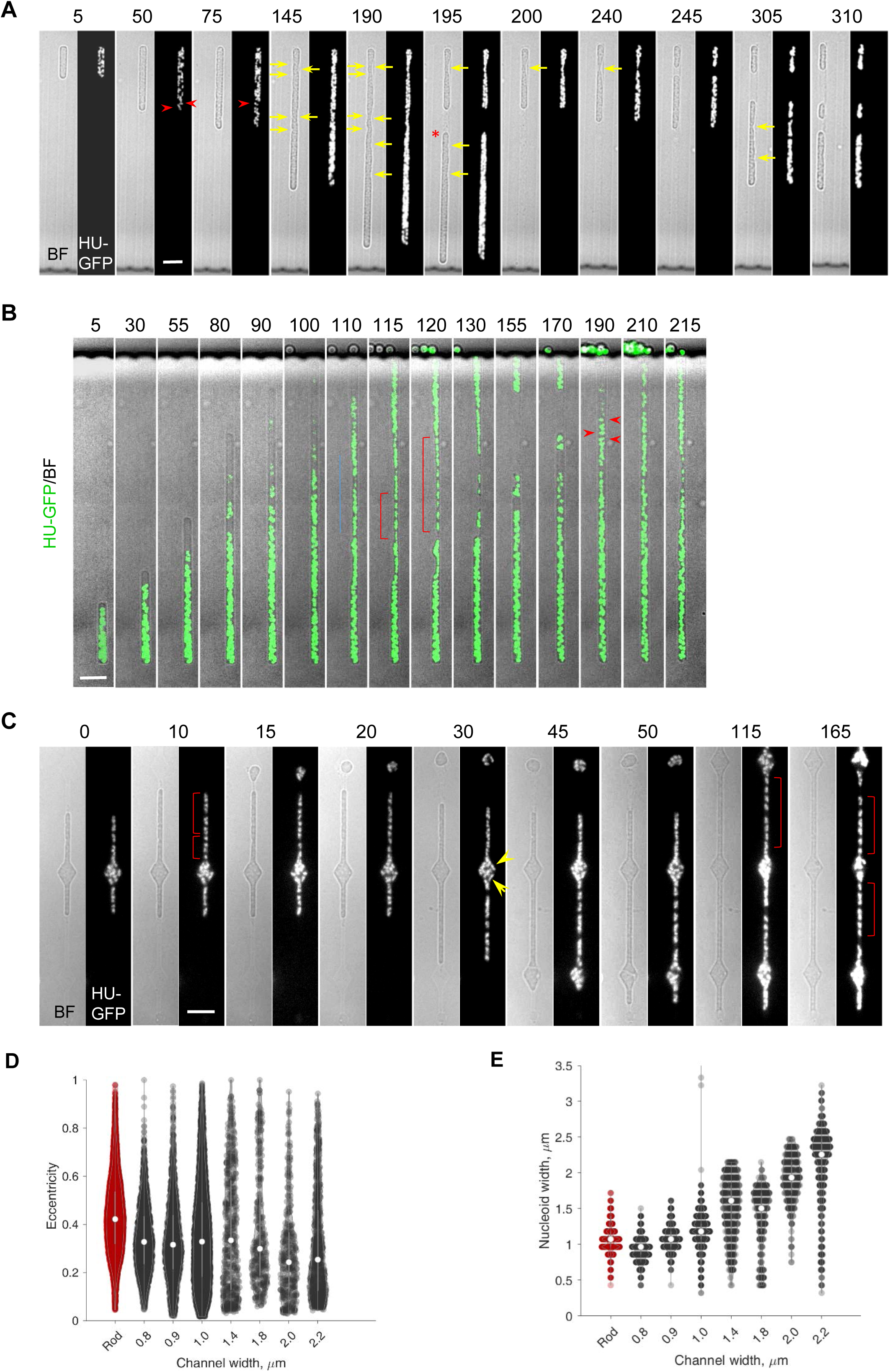
L-form division and chromosome distribution in microfluidic channels of various widths. Related to Figure 3. (A) In wide channels cell division occurred more frequently. However, chromosomes are not well separated and are distributed irregularly. These are selected still frames from Movie 3. Each time frame shows bright field images on the left and chromosomal DNA labelled with HU-GFP on the right. The cell shown was in a 2.0 µm wide channel (Chip No. 7). Red arrowheads: chromosomes lying horizontally or perpendicularly. Yellow arrows point to regions of membrane constrictions. The daughter cell that escaped from the channel is marked with a red star. Strain: 4739 (LR2 *ΩamyE*::*neo hbsU-gfp*). (B) Chromosomes became separated and more regularly distributed in the narrow parts of the wide cell (red brackets) where membrane constrictions persisted. The images are selected still frames from Movie 5, shown as the merge of the GFP image (green) and the bright field image (grey scale). The cell shown was in a 1.4 µm wide channel (Chip No. 6). Strain: 4739 (LR2 *ΩamyE*::*neo hbsU-gfp*). (C) In Chip No.33 which contained alternating narrow channels and diamond shapes, dis-organised chromosomes in the diamond parts became regularly distributed in the straight and narrow channels. These are selected still frames from Movie 6. Each time frame shows bright field images on the left and chromosomal DNA labelled with HU-GFP on the right. The channel width for the cell shown was 700 nm. Strain: 4741 (LR2 *ΩamyE*::*neo hbsU-gfp aprE::P_rpsD_-mcherry spc)*. Yellow arrowheads indicates two nucleoids in different orientations in the diamond region. Brackets in B & C indicate regions where chromosomes appear regularly distributed. Scale bars, 5 µm. (D) & (E): Nucleoid circularity (D) and width (E) for walled cells (red; Rod) and L-forms (black) in different channel widths. Violin plots: Circles indicate median, bars indicate upper and lower quartile. Scatter plots: Circles indicate median, error bars indicated 95 % confidence interval from bootstrapping. *n=17766* time-lapse observations of nucleoids descended from 45 mother cells in separate agarose channels.

**Figure S4:**
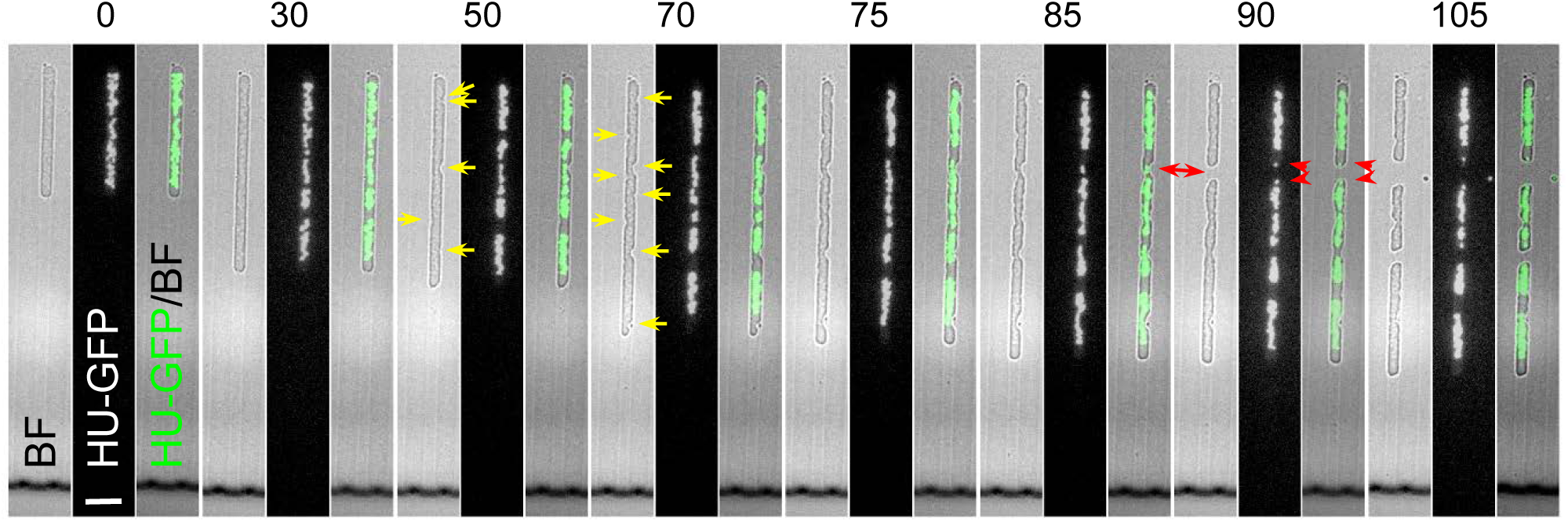
Bisection of chromosomes occurs occasionally in L-forms. Related to Figure 4. A chromosome appeared to have been bisected by division, with two small lobes of DNA (arrowheads in Frame 19) retained at the extreme ends of the cell where division has occurred (red arrows in Frames 85 and 90 min). These are selected still frames from Movie 8. The cell shown was in a 1.8 µm wide channel (Chip No. 7). Each time frame shows bright field images on the left, chromosomal DNA labelled with HU-GFP in the middle, and a merge of the two on the right (GFP in green). Yellow arrows point to regions of membrane constrictions. 5 mins between frames. Strain: 4739 (LR2 *ΩamyE*::*neo hbsU-gfp).* Scale bars, 5 µm.

**Figure S5:**
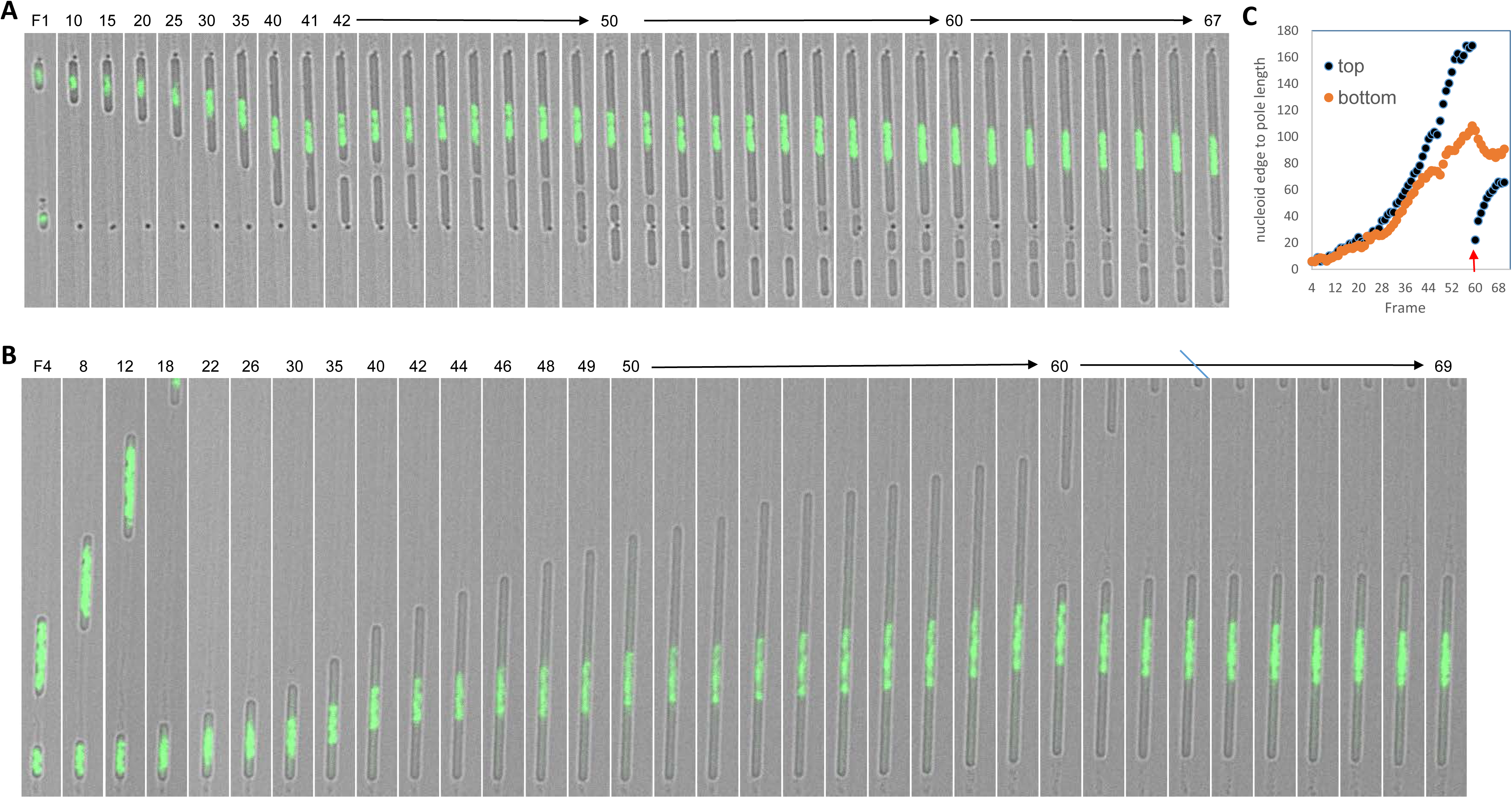
Re-centring of the single nucleoid in cells inhibited for DNA replication. (A) A fuller set of still images of a time-lapse experiment presented in Figure 6. The merge of the bright field image and the green fluorescence image of the chromosomal DNA labelled with HU-GFP (GFP in green) is shown. After division the nucleoid moved towards the distal pole to re-centre itself. L-forms of strain 4739 (LR2 *ΩamyE*::*neo hbsU-gfp)* were grown in the presence of the DNA replication inhibitor HB-EmAu in liquid culture and after introduction into a microfluidic device. 3 min per frame. Scale bars, 5 µm. (B, C) Another example of the single nucleoid re-centring after division.

**Figure S6:**
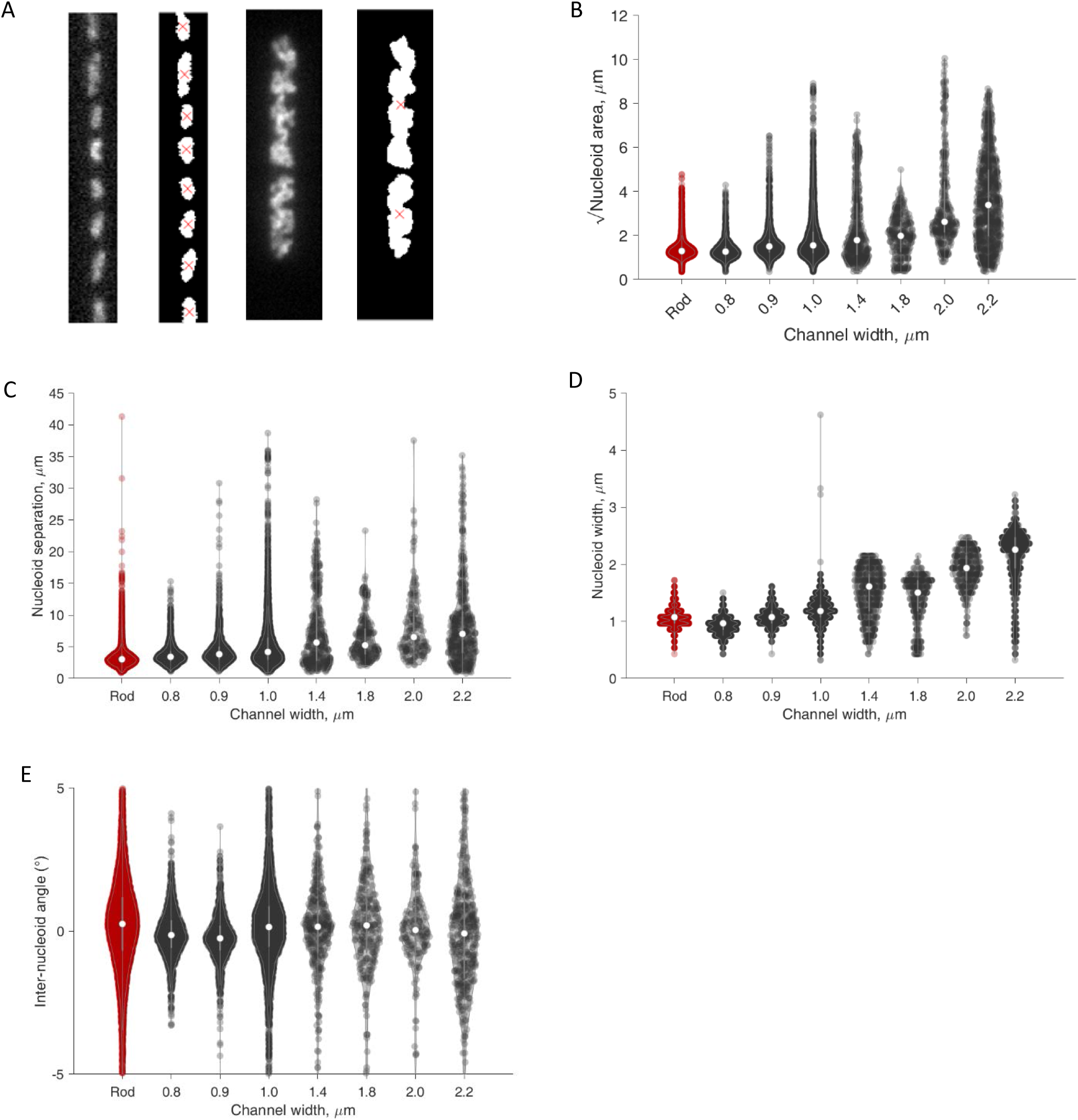
Further quantitative analysis of nucleoid shape. (A) Exemplar nucleoid segmentations for walled cells (left) and L-form cells in a 2.2 μm wide channel (right). (B-D) Violin plots showing full data range including outliers and extrema for Figures 3F, H and S3D. (E) Distribution of inter-nucleoid angle for all cells. Low frequency long tail of angles greater than 5 degrees not shown in order to visualize average trends.

**Movie 1**: L-forms grow out of focus when unstrained in liquid medium. Related to Figure 1A. A time-lapse experiment showing growth and chromosomes of L-form *B. subtilis* when growing unstrained in liquid medium in a glass-bottomed dish at 30°C. Chromosomes (green, merged with bright field images) soon became difficult to observe as the cells grew and divided in multiple directions. Strain: 4740 (LR2 *Pspac-dnaA ΩamyE*::*neo hbsU-gfp*). Phase contrast and the corresponding GFP images, which were overlaid, were acquired automatically every 5 min.

**Movie 2**: Some un-constrained L-form cells growing in the gutters of microfluidic systems remain in good focus. Related to Figure 1B. Time-lapse series with an agarose-based microfluidic system showing the growth of L-forms of strain 4745 (RM121 *ΩamyE*::*hbsU-mCherry rpoC-gfp)* in the gutter, from which the panels in Figure 1B were obtained. Phase contrast (left panel) and the corresponding HU-GFP (middle panel) images were acquired automatically every 5 min. Overlay of the phase contrast and the corresponding HU-GFP images is shown on the right.

**Movie 3**: L-form division and chromosome distribution in wide microfluidic channels (I). Related to Figure 3B. A time-lapse series showing cell division events and escaping of daughter cells of L-forms growing in wide channels. Chromosomes are not well separated and are distributed irregularly. The cell shown was in a 2.0 µm wide channel (Chip No. 7). Bright field images are shown on the left and the corresponding HU-GFP images showing the nucleoids are on the right. Strain: 4739 (LR2 *ΩamyE*::*neo hbsU-gfp*). 5 min between frames.

**Movie 4**: Another example of regular chromosome segregation in microfluidic channels. Time-lapse Series showing L-form cells of strain 4739 (LR2 *ΩamyE*::*neo hbsU-gfp*) growing in microfluidic channels (Chip No. 2; channel widths 0.8, 0.9 and 1.0 µm) at 32°C. Chromosomes can be seen segregating relatively regularly as the cells grew. Images were captured every 5 min. Bright field images are shown on the left and the corresponding HU-GFP images showing the nucleoids are on the right. Related to Figure 3A and Figure S2.

**Movie 5**: L-form division and chromosome distribution in wide microfluidic channels (II). Related to Figure S3B. A time-lapse series of merged HU-GFP (green) and bright field (grey scale) images showing irregularly distributed chromosomes in L-forms growing in wide channels becoming separated and more regularly distributed in the narrow parts of the cell where membrane constrictions persisted. The cell shown was in a 1.4 µm wide channel (Chip No. 6). Strain: 4739 (LR2 *ΩamyE*::*neo hbsU-gfp*). 5 min between frames.

**Movie 6**: L-form division and chromosome distribution in mixed shaped microfluidic device. Related to Figure 3E & S3C. A time-lapse series of HU-GFP (bottom) and bright field (top) images showing chromosome distribution in Chip No.33, which contained alternating narrow channels and diamond shapes. Dis-organised chromosomes in the diamond parts became regularly distributed in the straight and narrow channels. Strain: 4741 (LR2 *ΩamyE*::*neo hbsU-gfp aprE::P_rpsD_-mcherry spc)*. 5 min per frame.

**Movie 7**: Bisection of chromosomes in L-forms growing in wide channels. Related to Figure 4. Chromosomes (green, overlaid with the bright field images shown in grey scale) can be seen passing through areas of invagination, probably prevented division. A small cell with little amount of DNA (Frames 27 onwards) appeared not growing, probably because its chromosome is incomplete. The cell shown was in a 1.8 µm wide channel (Chip No. 7). 5 min between frames. Strain: 4739 (LR2 *ΩamyE*::*neo hbsU-gfp)*.

**Movie 8** Another example of chromosome bisection in L-forms growing in wide channels. Related to Figure S4. A chromosome (green, overlaid with the bright field images shown in grey scale) appeared to have been bisected by division, generating two small lobes of DNA (Frame 19) that retained at the extreme ends of the cell where division had occurred. The cell shown was in a 1.8 µm wide channel (Chip No. 7). Strain: 4739 (LR2 *ΩamyE*::*neo hbsU-gfp)*.

**Movie 9**: DNA-less ‘beads’ produced by L-form in narrow channels under normal growth conditions. Related to Figure 5A. The time-lapse series shows cells in the gutter grew into the narrow channels with the mass of the nucleoid excluded from entry, generating strings of DNA-less beads in narrow channels. Each frame shows chromosomes in green overlaid with the corresponding bright field image. Strain: 4739 (LR2 *ΩamyE*::*neo hbsU-gfp).* 5 min between frames.

**Movie 10:** Small DNA-free cells generated by divisions in DNA-free region of cells in wide channels. Related to Figure 5C. A cell, appeared to be defective in chromosome replication (for unknown reason), produced many small DNA-free daughter cells of various sizes. The cell shown was in a 2.2 µm wide channel (Chip No. 7). Each frame shows bright field images on the left and chromosomal DNA labelled with HU-mCherry on the right. 5 min between frames. Strain: 4739 (LR2 *ΩamyE*::*neo hbsU-gfp)*.

**Movie 11:** DNA-less cells produced in narrow channels by L-forms inhibited for DNA replication. The time-lapse series shows formation of DNA-less ‘beads on a string’. The replication inhibitor HB-EmAu was present in the medium throughout the time-lapse experiment. Strain: 4739 (LR2 *ΩamyE*::*neo hbsU-gfp).* 3 min between frames. Related to Figure 5D.

**Movie 12:** Large DNA-free cells divided into smaller cells. Related to Figure 5E. The time-lapse series shows formation of large DNA-less cells in narrow channels by L-forms inhibited for DNA replication, which divided further into smaller DNA-less daughters. The replication inhibitor HB-EmAu was present in the medium throughout the time lapse experiment. Strain: 4739 (LR2 *ΩamyE*::*neo hbsU-gfp).* 3 min between frames.

**Movie 13:** Re-centring of the single nucleoid after division. Time lapse of the experiment shown in Figure 6. The bright field image and the green fluorescence image of the chromosomal DNA labelled with HU-GFP (GFP in green) are merged. After division the asymmetrically located nucleoid moved towards the distal pole to re-centre itself. L-forms of strain 4739 (LR2 *ΩamyE*::*neo hbsU-gfp)* were grown in the presence of the DNA replication inhibitor HB-EmAu in liquid culture and after introduction into a microfluidic device. 3 min per frame.

**Table S1.**
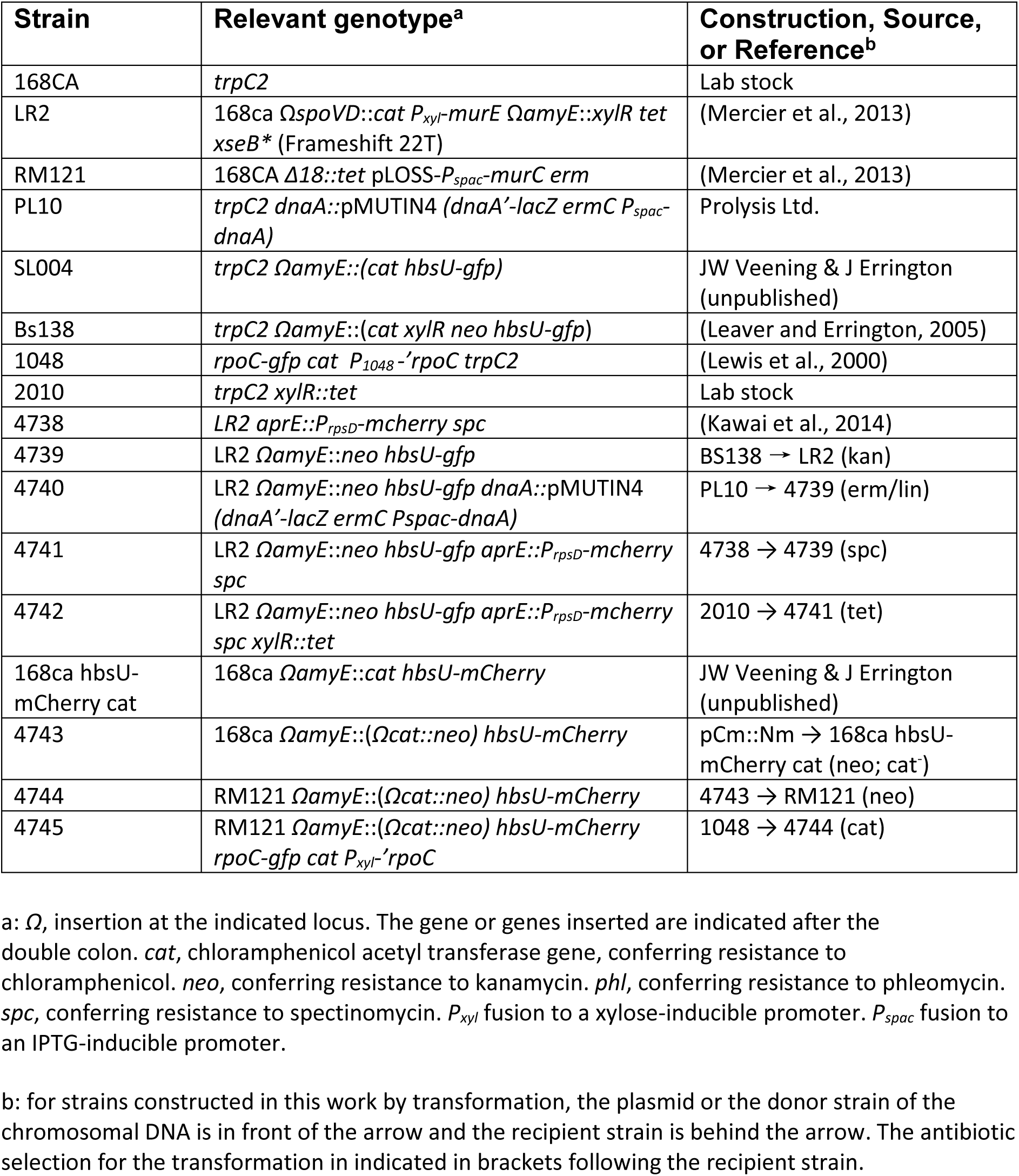
Bacterial strains and plasmid used in this study

